# Epigenetic Small-Molecule Screen for Inhibition and Reversal of Acinar Ductal Metaplasia in Mouse Pancreatic Organoids

**DOI:** 10.1101/2023.11.27.567685

**Authors:** Kalina R. Atanasova, Corey M. Perkins, Ranjala Ratnayake, Jinmai Jiang, Qi-Yin Chen, Thomas D. Schmittgen, Hendrik Luesch

**Affiliations:** Department of Medicinal Chemistry, College of Pharmacy, University of Florida, Gainesville, FL, USA; Center for Natural Products, Drug Discovery and Development, College of Pharmacy, University of Florida, Gainesville, FL, USA; Department of Pharmaceutics, College of Pharmacy, University of Florida, Gainesville, FL, USA

**Keywords:** pancreatic cancer, acinar ductal metaplasia, epigenetics, drug screening, organoids

## Abstract

**Background:** Acinar ductal metaplasia (ADM) is among the earliest initiating events in pancreatic ductal adenocarcinoma (PDAC) development.

**Methods:** We developed a novel morphology-based screen using organoids from wildtype and p48^Cre/+^ (Cre) mice to discover epigenetic modulators that inhibit or reverse pancreatic ADM more effectively than the broad-spectrum HDAC inhibitor trichostatin A (TSA).

**Results:** Of the 144 compounds screened, nine hits and two additional natural product HDAC inhibitors were validated by dose-response analysis. The class I HDAC inhibitors apicidin and FK228, and the histone methyltransferase inhibitor chaetocin demonstrated pronounced ADM inhibition and reversal without inducing significant cytotoxicity at 1 µM. Thioester prodrug class I HDAC inhibitor largazole attenuated ADM while its disulfide homodimer was effective in both ADM inhibition and reversal. Prioritized compounds were validated for ADM reversal in p48^Cre/+^;LSL-Kras^G12D/+^ (KC) mouse organoids using both morphological and molecular endpoints. Molecular index analysis of ADM reversal in KC mouse organoids demonstrated improved activity compared to TSA. Improved prodrug stability translated into a stronger phenotypic and molecular response. RNA-sequencing indicated that angiotensinogen was the top inhibited pathway during ADM reversal.

**Conclusion:** Our findings demonstrate a unique epigenetic mechanism and suggest that the phenotypic screen developed here may be applied to discover potential treatments for PDAC.

## 1 Introduction

Pancreatic ductal adenocarcinoma (PDAC) is among the most lethal cancers worldwide [1] with the majority of cases being discovered after metastasis has occurred [2; 3; 4]. The dire outcomes are in great part due to lack of effective treatments and poor methods of early detection. Understanding the early stages of PDAC development is critical to improve early detection and treatment strategies. Studies in genetically modified mouse models reveal an underlying role for pancreatic acinar cells in PDAC development [5; 6; 7; 8]. One of the earliest known initiating events for PDAC is the process of acinar ductal metaplasia (ADM) [9; 10]. During ADM, acini transdifferentiate into duct-like structures with reduced expression of acinar markers such as amylase (AMY2A) or carboxypeptidase (CPA2) and increased ductal markers such as cytokeratin 19 (KRT19). Following inflammatory or other insults, ADM is a natural process that occurs to protect acinar cells from further enzymatic damage. In mouse models of PDAC, ADM is achieved by restricted expression of *Kras* in acinar cells, [8; 11] experimental pancreatitis, [12] TGF-α [13] or other factors that activate EGFR [10]. ADM is believed to be irreversible with respect to mutant *Kras* [14; 15; 16].

The current FDA-approved therapies for PDAC include gemcitabine in combination with platinum agents or the FOLFIRINOX drug combination [17]. Many drugs in clinical trials for treating PDAC such as romidepsin (FK228), vorinostat (SAHA) or curcumin [18; 19] rely on epigenetic mechanisms to induce cell death of tumor cells. However, despite initial successful effects in clinical trials, many of these epigenetic drug trials have been terminated due to high toxicity [18].

Using primary cultures of mouse pancreas at different days of embryonic development, treatment with the histone deacetylase (HDAC) inhibitor trichostatin A (TSA) reduced acinar differentiation and promoted ductal differentiation [20]. Treatment of cultured acinar cells with TSA reduced caerulein-induced trypsin activation [21]. Using the caerulein-induced pancreatic injury model in mice, treatment with the HDAC inhibitor valproic acid delayed recovery of the pancreas, reduced acinar cell proliferation, maintaining ADM and thus delaying acinar redifferentiation [22]. HDAC expression, in particular class I HDAC, was upregulated during caerulein-induced pancreatitis in mice and inhibition of class I HDAC with MS-275 reduced ADM both in vitro and in vivo [23]. We recently showed that TSA or the STAT3 inhibitor LLL12B inhibited ADM formation in a mouse acinar organoid 3D culture model [24]. Moreover, TSA exposure following completion of ADM induced phenotypic and gene expression changes reminiscent of ADM reversal in organoids from mice carrying the *Kras^G12D^* mutation [24]. *KRAS^G12D^* is among the most common mutations observed in PDAC patients and is considered to be one of the main drivers of ADM and progression to PDAC [25; 26; 27; 28; 29].

Given the need for new, effective therapeutics with minimal toxicity, we developed a novel medium-throughput screen to discover epigenetic modulating compounds that inhibit and/or reverse ADM. We utilized the mouse acinar organoid 3D-culture model and monitored ADM or its reversal using high-content organoid morphology as a readout. We screened the Cayman’s small-molecule epigenetic modulator library (ESL) which contains 144 compounds with the goal to identify important regulators of ADM with therapeutic potential. A large component of the ESL library (43 compounds) consists of HDAC inhibitors with broad or narrow molecular target and isoform range. Of these, 34 HDAC inhibitors were Zn^2+^ - dependent (targeting HDACs class I, II or IV) with differential class or isoform selectivity (class I vs II vs I/II) selectivity and 9 were NAD^+^-dependent (targeting class III HDACs).

During the initial screen, we identified nine epigenetic modulating compounds, two of which were class I HDAC inhibitors, that induced phenotypic changes with high mechanism-dependent selectivity and low cytotoxicity. The effects of these nine compounds and additional class I-selective HDAC inhibitor prodrugs, largazole (thioester), largazole homodimer (disulfide), and the functionally similar natural product FK228 were further validated in dose-response studies using the same model. The prioritized compounds were effective at re-expression of acinar genes (*Amy2a, Cpa2*) and suppressing ductal genes (*Krt19*, *Sox9*) in the *Kras^G12D^* mouse organoid model demonstrating that ADM is reversible even in the context of mutant *Kras*.

## 2 Materials and Methods

### 2.1 Mice

C57BL/6J (wild type) mice were used in ADM inhibition assays. p48^Cre/+^ mice (Cre-mice) were bred to LSL-Kras^G12D/+^ to produce p48^Cre/+^; LSL-Kras^G12D/+^ mice (KC mice). Genotyping for the presence of the transgene was performed by Transnetyx (Cordova, TN).

### 2.2 Mouse acinar ductal metaplasia culture

Acinar cells were isolated as previously reported [24] from the pancreas of 6–8-week-old wild type (ADM inhibition assays), Cre-mice (ADM reversal assays), or KC mice (ADM reversal assays). Following a 30- minute digestion with collagenase P, enzyme inactivation with fetal bovine serum containing Hanks’ balanced salt solution, and a series of centrifugation steps, cells were passed through a series of strainers (500, 300, 200 µm). The resulting pellet was resuspended in media to the desired density to yield 50 organoids/well with growth factor reduced Matrigel (Corning). The mixture was maintained on ice during the pipetting steps. Twenty µL of the mixture was pipetted into a 384-well CellCarrier Ultra plates (PerkinElmer), followed by 30 minutes of incubation to solidify the mixture and then 40 µL of warm media was added. The organoids seeded in Matrigel were left overnight to acclimate and then subjected to treatment or left to further develop ADM before treatment.

### 2.3 Compound screen, reagents, treatment and sample processing

#### 2.3.1 ADM Inhibition and Reversal Experimental Design

In ADM inhibition experiments wild type mouse organoids were imaged live after overnight acclimation using the Operetta High Throughput Screen imaging system (PerkinElmer) using 20× objective, brightfield filter and 15 fields of view/well covering ∼80% of the well area. The wells were immediately treated using a JANUS liquid handling system (PerkinElmer) and a 200 nL pin tool. In ADM reversal experiments p48^Cre/+^ (screen and validation experiments) or KC-mice organoids (follow up experiments) were left to incubate at 37°C, 5% CO_2_ for additional 72 or 48 hours, respectively, until ducts formed, then fresh media was exchanged and the organoids were imaged and treated in same manner as for ADM inhibition experiments.

#### 2.3.2 Epigenetic Compound Treatment

For screening experiments, organoids were treated with the Cayman small-molecule epigenetic library (Cayman Chemicals, #11076) at final concentration of 1 µM. For validation studies, organoids were treated with a relevant concentration range of 3-fold dilutions of the selected compounds. Vehicle control wells (0.5% DMSO, n=4) were included on every plate. TSA (Sigma-Aldrich, #T8552) was used as a positive control based on previous findings [24] and was tested in a 3-fold dose range from 32 nM to 10 µM (quadruplicate wells per dose). FK228 (romidepsin) was purchased from Sigma-Aldrich, (#SML1175).

#### 2.3.3 Viability Staining and Imaging

Post treatment plates were incubated at 37°C, 5% CO_2_ for 72 hours and the images were visualized for duct/cluster-like morphology. Media was removed and organoids in Matrigel were washed once with 100 µL/well of Dulbecco’s phosphate buffered saline (DPBS, Corning), followed by staining with Calcein AM (Sigma) in DPBS at final concentration of 4 µM for 1.5 hours at 37°C in the dark. After staining, organoids were washed again with 100 µL/well of DPBS and left in fresh DPBS for live imaging. Organoids were imaged on Operetta High-Throughput Screen imaging system using 20× objective, brightfield and green fluorescence filters at the same 15 fields/well that were imaged before the treatment.

#### 2.3.4 Duct/Cluster-Like Counting

Following 72 h of treatment, total and live (Calcein AM positive) duct and cluster counts were compared to the vehicle control treated wells to determine cytotoxicity. Brightfield (both before treatment and 72 h post treatment) and green-fluorescence filter (at 72 h post treatment) images were analyzed by identification of the primary objects in stitched images of the 15 fields/well using a pixel classification of background/objects-based procedure in Ilastik 1.3.2 [30] and a custom-built pipeline in CellProfiler 3.1.8 software [31], followed by object quantification, and viability quantification (using the green channel, calcein AM staining images). Objects were classified as either duct-like or cluster-like using the machine learning object classification software CellProfiler Analyst 3.0.3 [32] based on the properties/features collected from the CellProfiler software. All stitched images were also visually assessed to confirm observations from the machine learning components and the robustness of the pipeline/classification process, as well as to detect morphological changes that did not fall within the duct-like/cluster-like classification, e.g. duct size changes and other morphological changes. ADM at 72 h post treatment was expressed as the percent ratio of viable ducts and viable acinar clusters from all live objects for both the ADM inhibition and the ADM reversal screens. Analysis of experiments carried out for the selected few hits done in KC mice was performed manually by visualization due to the developed pipeline not being able to recognize texture and morphological differences specific for the cyst-like organoids compared to the acinar clusters.

### 2.4 RNA isolation and quantitative gene expression

To process the 384 well plates for RT-qPCR, we used a modification of our method to remove Matrigel from the cell mixture [33]. Media was removed by inverting the plate onto paper towels to blot off the media. The plate was then placed on ice for 10 minutes to liquefy the Matrigel. Forty µL of ice-cold PBS was added to each well of the plate on ice. The contents of 16 wells were then combined per treatment by removing 60 µl from each well and combined into a 2 ml centrifuge tube on ice. An additional 1 mL of cold PBS was added to the tube and resuspended. The tubes were centrifuged at 1000 × G for 5 mins at 4° C. The supernatant was removed by pipetting and then resuspended in 2 ml of cold PBS, followed by an additional centrifugation step at 1200 × G for 5 mins at 4 °C. Supernatant was removed by pipetting and the cell pellet was resuspended in 700 µL of TRIzol reagent. Total RNA was isolated using the miRNeasy protocol (Qiagen). Sixty ng of total RNA was converted to cDNA in a 20 µL RT reaction using random primers and MMLV reverse transcriptase (Thermo). qPCR was performed using the QuantStudio™ 7 Flex Real-Time PCR System (Thermo). Data are presented using the 2^-ΔΔCT^ method [34] relative to vehicle control and normalized to 18S rRNA. Primer sequences have been previously published [24].

### 2.5 RNA sequencing and pathway enrichment analysis

Illumina RNA library construction was performed at the Interdisciplinary Center for Biotechnology Research (ICBR) Gene Expression Core, University of Florida (UF). RNA quantitation was done on a NanoDrop Spectrophotometer (NanoDrop Technologies, Inc.), and sample quality was assessed using the Agilent 2100 Bioanalyzer (Agilent Technologies, Inc). SMART-Seq V4 ultra low input RNA kit were used for RNAseq library construction according to the user manual. Illumina sequencing libraries were generated with 125 pg of cDNA using Illumina Nextera DNA Sample Preparation Kit (Cat#: FC-131-1024) according to manufacturer’s instructions. The libraries were pooled in equal molar concentration. Normalized libraries were submitted to the “Free Adapter Blocking Reagent” protocol (FAB, Cat# 20024145) in order to minimize the presence of adaptor-dimers and index hopping rates. The library pool was diluted to 0.8 nM and sequenced on one S4 flow cell lane (2×150 cycles) of the Illumina NovaSeq6000 using NovaSeq Control Software v1.6. Sample sequencing was performed at the ICBR NextGen Sequencing (https://biotech.ufl.edu/next-gen-dna/, RRID:SCR_019152). Additional details on library generation and sequencing are given in the supplementary materials and methods.

Reads acquired from the Illumina NovaSeq 6000 platform were cleaned up with the cutadapt program [35] to trim the sequencing adaptors and low-quality bases with a quality phred-like score <20. Reads <40 bases were excluded from RNA-seq analysis. The genome of *Mus musculus* (version GRC38, mm10) from the Ensembl database was used as the reference sequences for RNA-seq analysis and cleaned reads were mapped to the reference sequences using the read mapper of the STAR package (Spliced Transcripts Alignment to a Reference, v2.7.9a) [36]. The mapping results were processed with the HTSeq (High-Throughput Sequence Analysis in Python, v0.11.2) [37], samtools, and scripts developed in house at ICBR to remove potential PCR duplicates and count uniquely mapped reads for gene expression analysis. Outliers were detected using PCA analysis and volcano plot analysis based on all identified genes using R-package (v4.1.3). The gene express levels were analyzed by a DESeq2-based R pipeline.

Gene expression levels of approximately 4,000 differentially expressed genes compared to vehicle control at 72 h post treatment were compared using Ingenuity Pathway Analysis software (QIAGEN Inc., https://digitalinsights.qiagen.com/IPA) and hits were considered as significant when they passed the following cut-offs: fold change difference greater than 1.5 and a false discovery rate (FDR) p value of less than 0.05. The z-score was calculated dependent upon the fold change and FDR requirements, resulting in the identification of the top upstream and downstream effectors following reversal treatment.

### 2.6 Statistical analysis and selection of compounds for validation

Percent live ducts for each treatment was compared to vehicle control and p values were calculated using two-tailed Student’s *t*-test with unequal variances. The Z’-score for evaluating the optimization quality of the assay was calculated using negative control (vehicle only: DMSO, 0.5%) and positive control at concentration inducing inhibition or reversal of ADM in over 80% of all organoids in the well (TSA, 10 µM) [38]. Significance was considered at p ≤ 0.05. Compounds that showed significant decrease in % live ducts compared to vehicle control, while showing over 50% total viable objects, were selected for further validation and analysis. The ADM Reversal Index (ADMRI) is defined as the mean fold change in expression (treated versus control) as determined by qRT-PCR of acinar genes (Amy2a, Cela and Cpa2) divided by the mean fold change in ductal gene (Krt19, Krt7 and Sox9) expression [24]. We annotated 2 sets of genes previously associated with acinar/ductal phenotype of pancreas or genes associated with onset or progression of PDAC, from the literature [39; 40]. Volcano plots were constructed from the changes in expression of 27 acinar and 23 ductal/PDAC genes following ADM reversal as previously described [41]. A list of the acinar and ductal/PDAC genes may be found in Table S1 [39; 40].

## 3 Results

### 3.1 Assay validation

Experimental efficiency was verified in both the inhibition (wildtype mouse organoids) and reversal (*p48^Cre^* (Cre) mouse organoids) screening modes using TSA as the positive control. A concentration-dependent relationship was observed for both inhibition and reversal effects while cell morphology showed the presence of viable acinar clusters rather than viable ducts as with the vehicle (0.5% DMSO) controls (Figure 1). The Z’ for both the ADM inhibition (Z’ = 0.61) and ADM reversal (Z’ = 0.56) was calculated using the positive control TSA. The values for Z’ was > 0.50 indicating that both assays were sufficiently optimized for high-throughput screening.

**Figure 1.**
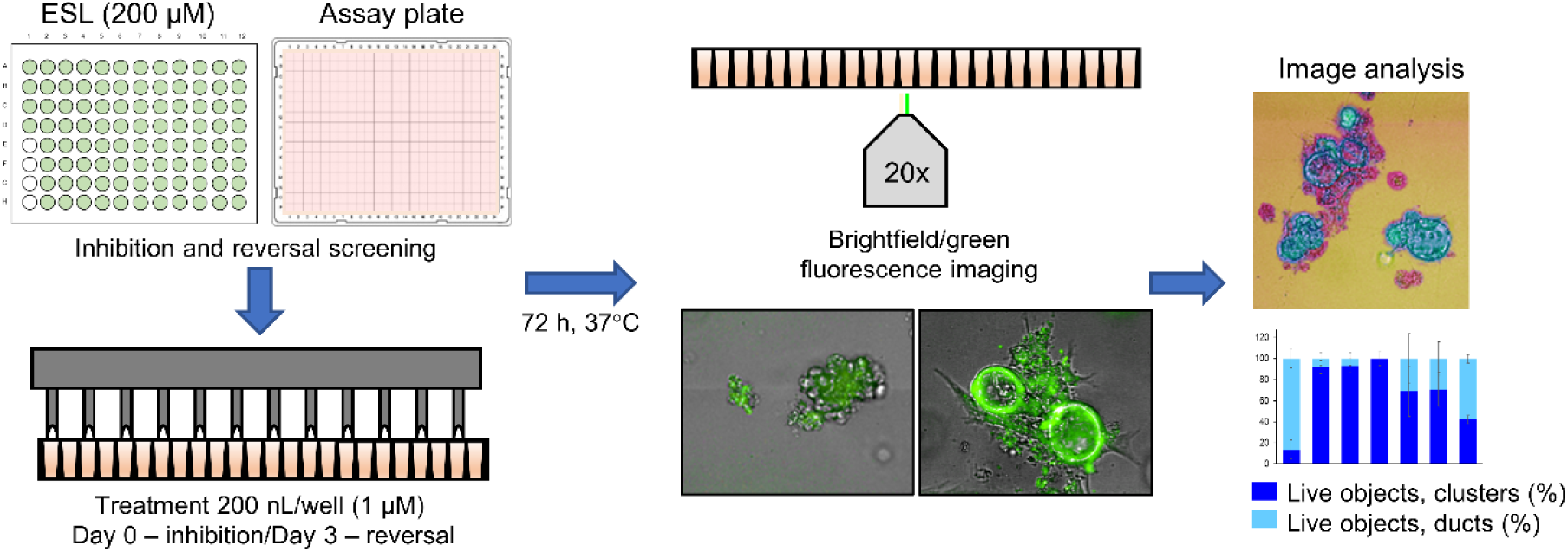
Distribution of viable ducts and acinar clusters in trichostatin A (TSA)-treated organoids. Percent duct/cluster distribution (± standard deviation) of live objects at 72 h post treatment with positive control TSA as a function of the concentration in ADM inhibition (wildtype mouse organoids, Z’ = 0.61) (A) and ADM reversal (Cre-mice organoids, Z’ = 0.56) (B) assays. Representative images at 1 µM TSA for ADM inhibition (C) and ADM reversal (D) assays. Scale bars represent 100 µm.

### 3.2 Epigenetic Small-Molecule Library Screen (ADM Inhibition)

The screen of 144 epigenetic modulating compounds for ADM inhibition was performed on wildtype mouse organoids. Schematic representation of the screening workflow is shown in Figure 2. Results from the inhibition screening assays were subjected to three criteria in order to select hits for further validation (Figure 3): *i*) compounds that showed a higher percentage of viable acinar clusters compared to the vehicle control (P<0.05), *ii*) compounds that showed a higher percentage of total clusters (viable and nonviable) compared to the vehicle control (P<0.05), and (*iii*) the passing compounds from *i*) and *ii*) produced ≤50% cytotoxicity 72 h post treatment as measured by calcein AM staining (for detailed results see Figures S1- S4).

**Figure 2.**
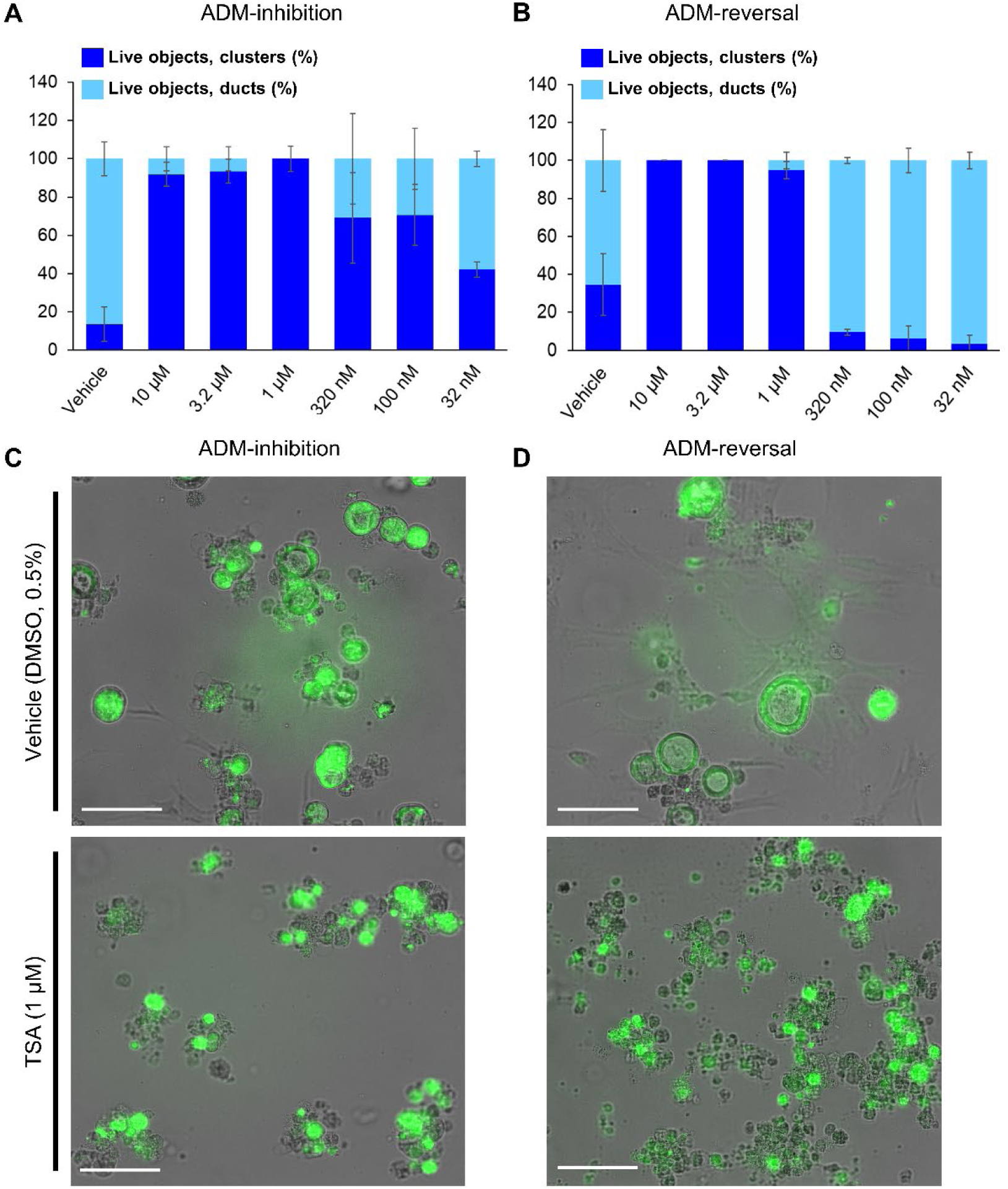
Schematic representation of the ADM screen. Seeded organoids originating from wildtype (WT) or p48^Cre/+^ (Cre) mice were treated with the Cayman small-molecule epigenetic modulator library at final concentration of 1 µM using an automated dispensing system and left to incubate for 72 h. Following incubation, organoids were stained using the calcein AM viability dye and imaged using a high throughput imaging system at 20× magnification to obtain high resolution images for further image analysis.

**Figure 3.**
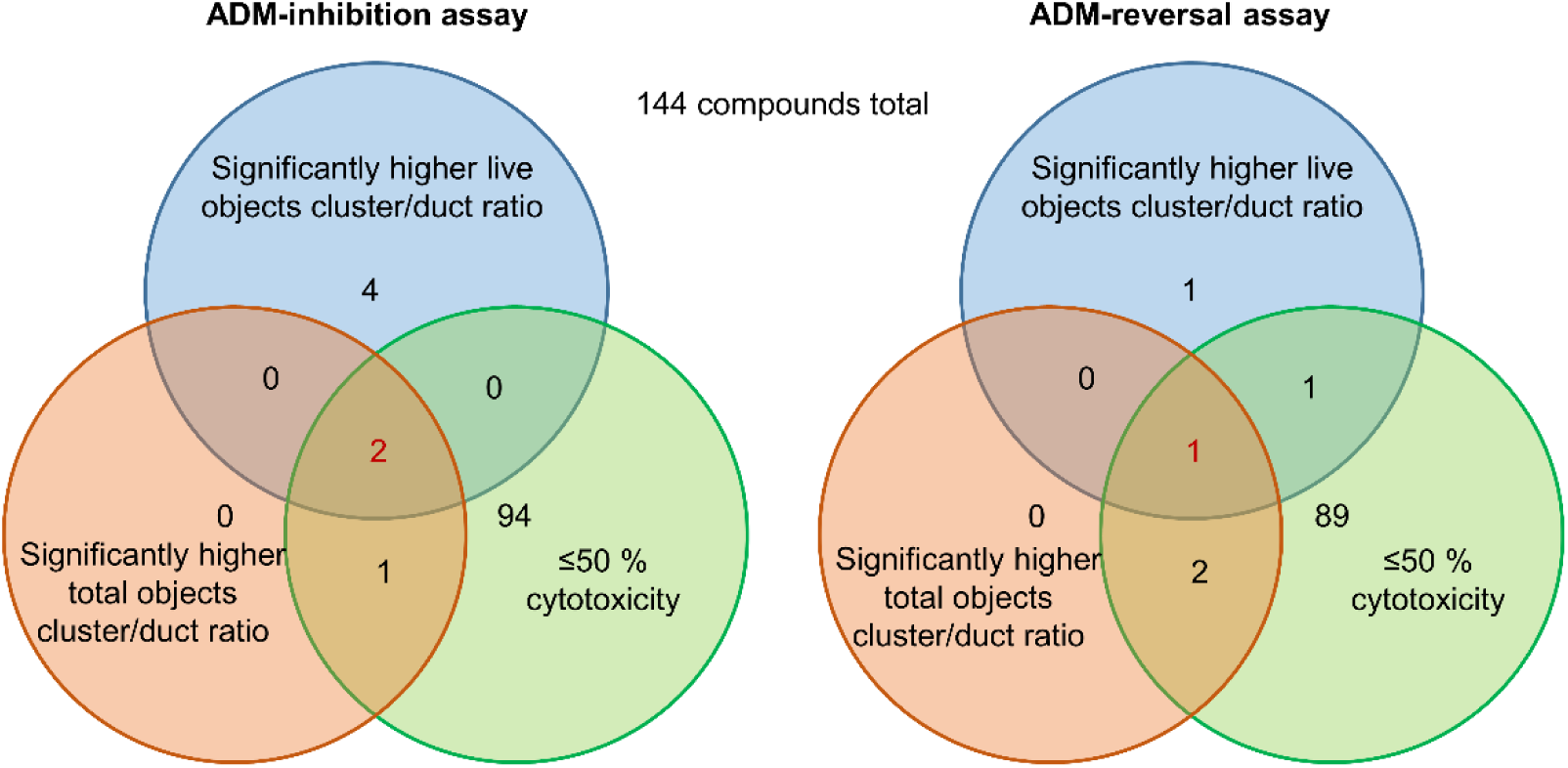
Selection criteria for compound validation. Summary of selection criteria applied to prioritize compounds for validation from ADM inhibition (wildtype mice organoids) and ADM reversal (Cre-mice organoids) screens. Cytotoxicity was evaluated by calcein AM staining. Compounds that overlapped in all three selection criteria (apicidin and chaetocin in ADM inhibition assay; chaetocin in ADM reversal assay) are shown in red and were prioritized for validation in dose response.

Of the 144 compounds screened for ADM inhibition, 6 compounds showed significantly higher percentages of live clusters at the tested concentration of 1 µM compared to the vehicle control (chaetocin, apicidin, IBET151, OTX015, 3-deazaneplanocin A and B32B3). Four of those compounds (IBET762, OTX015, 3- deazaneplanocin A and B32B3) resulted in >50% cytotoxicity based on the calcein AM staining (Figure 4, 5 and Figures S1 and S3). The two prioritized compounds from the inhibition screen are apicidin (Zn^2+^ dependent, class I HDAC inhibitor) and the histone methyltransferase (HMT) inhibitor chaetocin (Figures 4 and 5), fulfilling our criteria.

**Figure 4.**
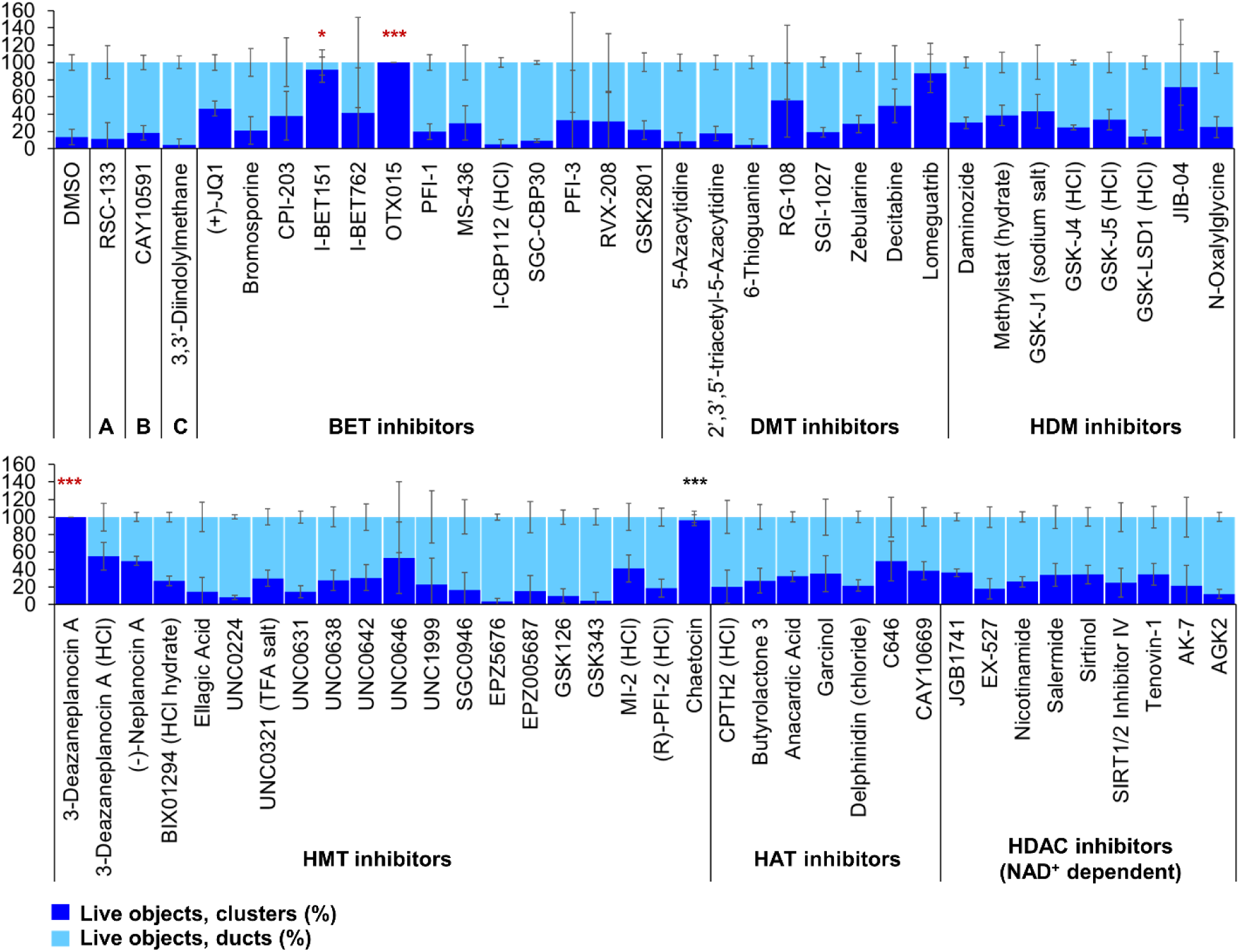
ADM inhibition from epigenetic compound library screen (compounds 1-68 grouped by target). The ADM inhibition assay screen was performed in duplicate using the Cayman ESL on wildtype mouse organoids. The mean percentage of viable ducts and acinar clusters (± standard deviation) 72 h post treatment. BET = bromodomain and extraterminal domain inhibitors; DMT = DNA methyltransferase inhibitors; HDM = histone demethylase inhibitors; HMT = histone methyltransferase inhibitors; HAT = histone acetyltransferase inhibitors; HDAC = histone deacetylase inhibitors (NAD^+^ dependent); HCl = hydrochloride; TFA salt = trifluoroacetic acid salt. P values were calculated using two-tailed Student’s t-test with unequal variances. Significance was accepted at p≤0.05 only when averages of clusters are higher than the respective vehicle control. * p value = 0.05 – 0.01; ** p value = 0.01 – 0.001; *** p value <0.001. Red asterisks denote compounds that showed significantly higher cluster/duct ratios of live objects but had overall less than 50% viability of total objects (false positive). Black asterisks denote compounds that showed significantly higher cluster/duct ratios of live objects and more than 50% of total objects were viable by calcein AM staining (true positive).

**Figure 5.**
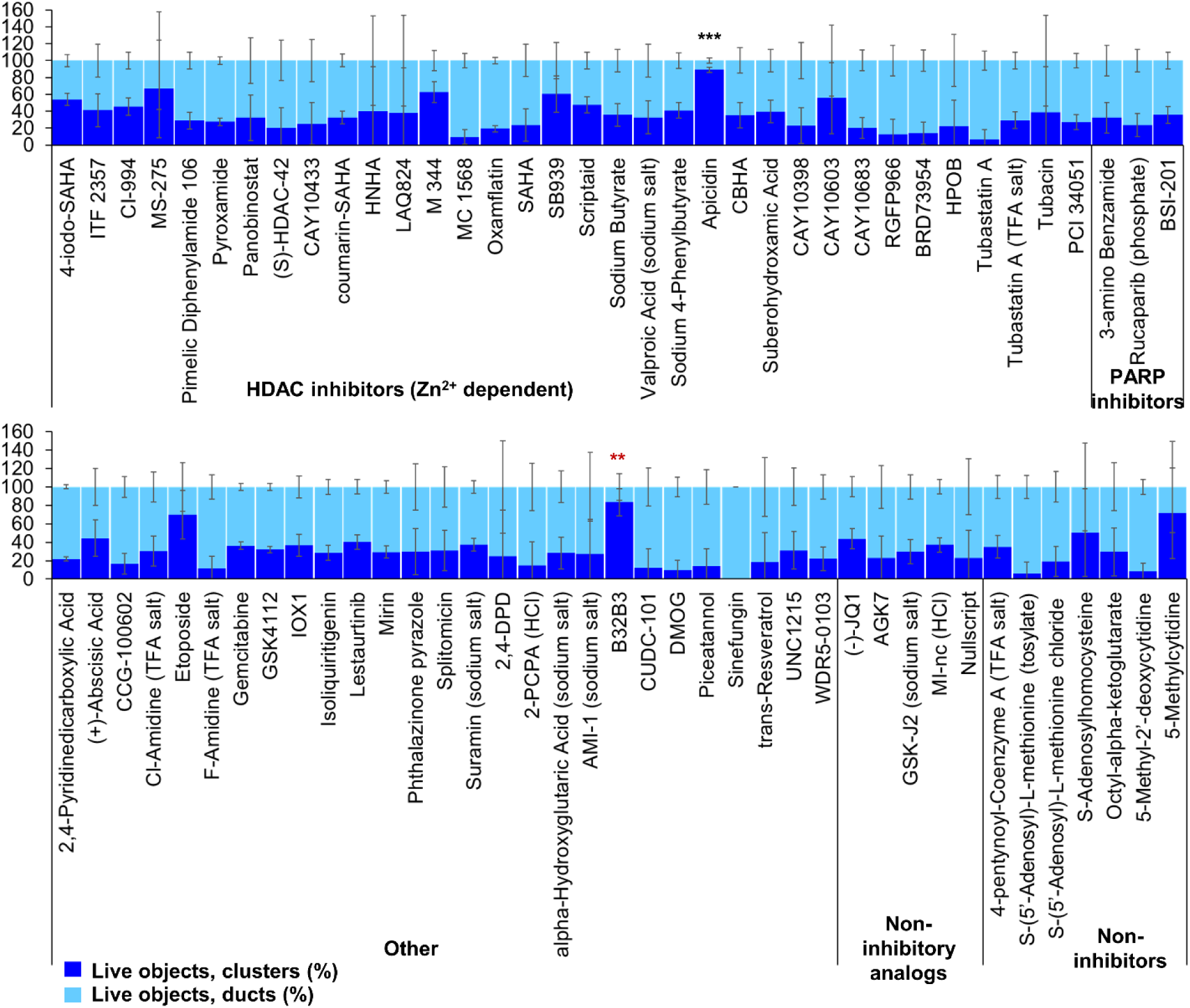
ADM inhibition from epigenetic compound library screen (compounds 69-144 grouped by target). The ADM inhibition assay screen was performed in duplicate using the Cayman ESL on wildtype mouse organoids. The mean percentage of viable ducts and acinar clusters (± standard deviation) 72 h post treatment. HDAC = histone deacetylase inhibitors (Zn^2+^ dependent); PARP = poly-ADP ribose polymerase inhibitors; HCl = hydrochloride; TFA salt = trifluoroacetic acid salt. P values were calculated using two-tailed Student’s *t*-test with unequal variances. Significance was accepted at p≤0.05 only when averages of clusters are higher than the respective vehicle control. * p value = 0.05 – 0.01; ** p value = 0.01 – 0.001; *** p value <0.001. Red asterisks denote compounds that showed significantly higher cluster/duct ratios of live objects but had overall less than 50% viability of total objects (false positive). Black asterisks denote compounds that showed significantly higher cluster/duct ratios of live objects and more than 50% of total objects were viable by calcein AM staining (true positive).

### 3.3 Epigenetic Small-Molecule Library Screen (ADM Reversal)

The screen of 144 epigenetic modulating compounds for ADM reversal was performed on Cre mouse organoids. The criteria to prioritize compounds from the screen were identical to those described for ADM inhibition. Only two compounds (chaetocin and apicidin) passed our rigorous selection criteria (Figures 6 and 7). The class II HDAC (HDAC6) inhibitor tubastatin A passed criteria *i* and *ii* however it induced > 50% cytotoxicity (Figures 6, 7, S2 and S4).

**Figure 6.**
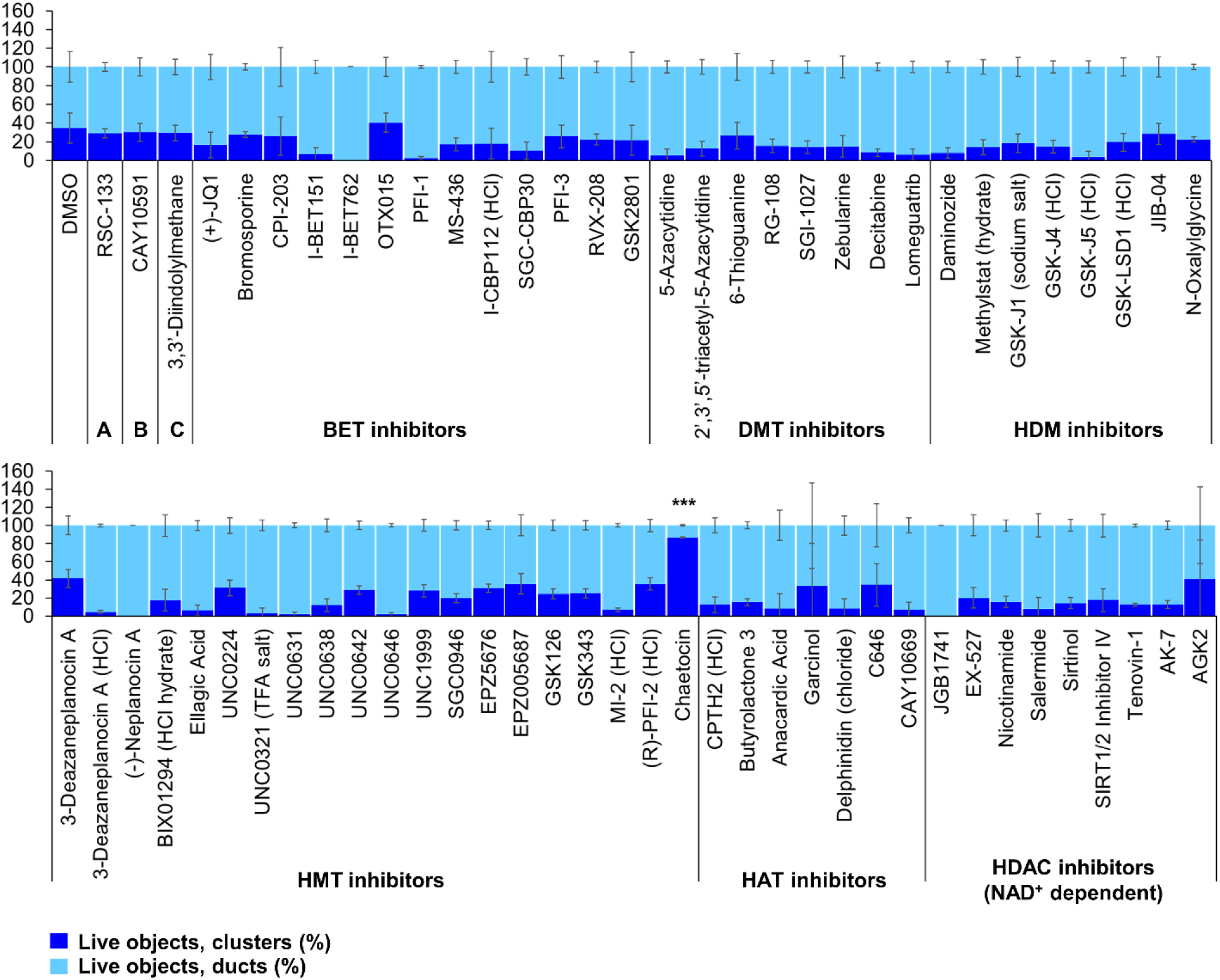
ADM reversal from epigenetic compound library screen (compounds 1-68 grouped by target). The ADM inhibition assay screen was performed in duplicate using the Cayman ESL on wildtype mouse organoids. The mean percentage of viable ducts and acinar clusters (± standard deviation) 72 h post treatment. BET = bromodomain and extraterminal domain inhibitors; DMT = DNA methyltransferase inhibitors; HDM = histone demethylase inhibitors; HMT = histone methyltransferase inhibitors; HAT = histone acetyltransferase inhibitors; HDAC = histone deacetylase inhibitors (NAD^+^ dependent); HCl = hydrochloride; TFA salt = trifluoroacetic acid salt. P values were calculated using two-tailed Student’s t-test with unequal variances. Significance was accepted at p≤0.05 only when averages of clusters are higher than the respective vehicle control. * p value = 0.05 – 0.01; ** p value = 0.01 – 0.001; *** p value <0.001. Red asterisks denote compounds that showed significantly higher cluster/duct ratios of live objects but had overall less than 50% viability of total objects (false positive). Black asterisks denote compounds that showed significantly higher cluster/duct ratios of live objects and more than 50% of total objects were viable by calcein AM staining (true positive).

**Figure 7.**
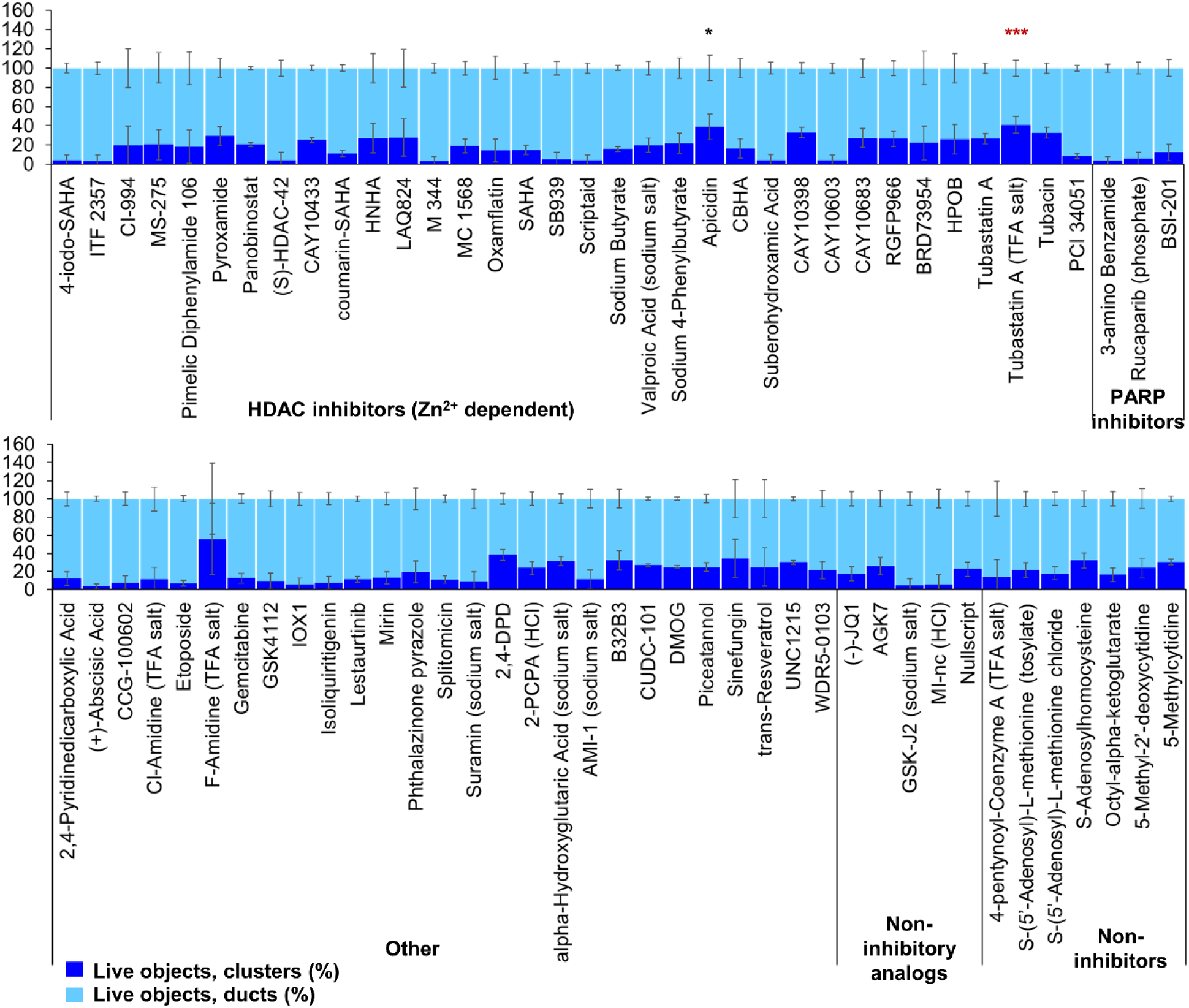
ADM reversal from epigenetic compound library screen (compounds 69-144 grouped by target). The ADM inhibition assay screen was performed in duplicate using the Cayman ESL on wildtype mouse organoids. The mean percentage of viable ducts and acinar clusters (± standard deviation) 72 h post treatment. HDAC = histone deacetylase inhibitors (Zn^2+^ dependent); PARP = poly-ADP ribose polymerase inhibitors; HCl = hydrochloride; TFA salt = trifluoroacetic acid salt. P values were calculated using two-tailed Student’s *t*-test with unequal variances. Significance was accepted at p≤0.05 only when averages of clusters are higher than the respective vehicle control. * p value = 0.05 – 0.01; ** p value = 0.01 – 0.001; *** p value <0.001. Red asterisks denote compounds that showed significantly higher cluster/duct ratios of live objects but had overall less than 50% viability of total objects (false positive). Black asterisks denote compounds that showed significantly higher cluster/duct ratios of live objects and more than 50% of total objects were viable by calcein AM staining (true positive).

In addition to the automated, quantitative data obtained from the screen, we visually observed the sets of images collected from each of the treatments for both inhibition and reversal to see if any compounds produced unique morphological characteristics that could not be detected by our pipeline. LAQ824 reduced duct size during inhibition (Figure 8). The BET inhibitors IBET762, IBET151, (+) JQI and PFI-1 produced notably enlarged ducts during ADM inhibition but not during ADM reversal (Figure 8). The findings for morphology are summarized in Table 1.

**Figure 8.**
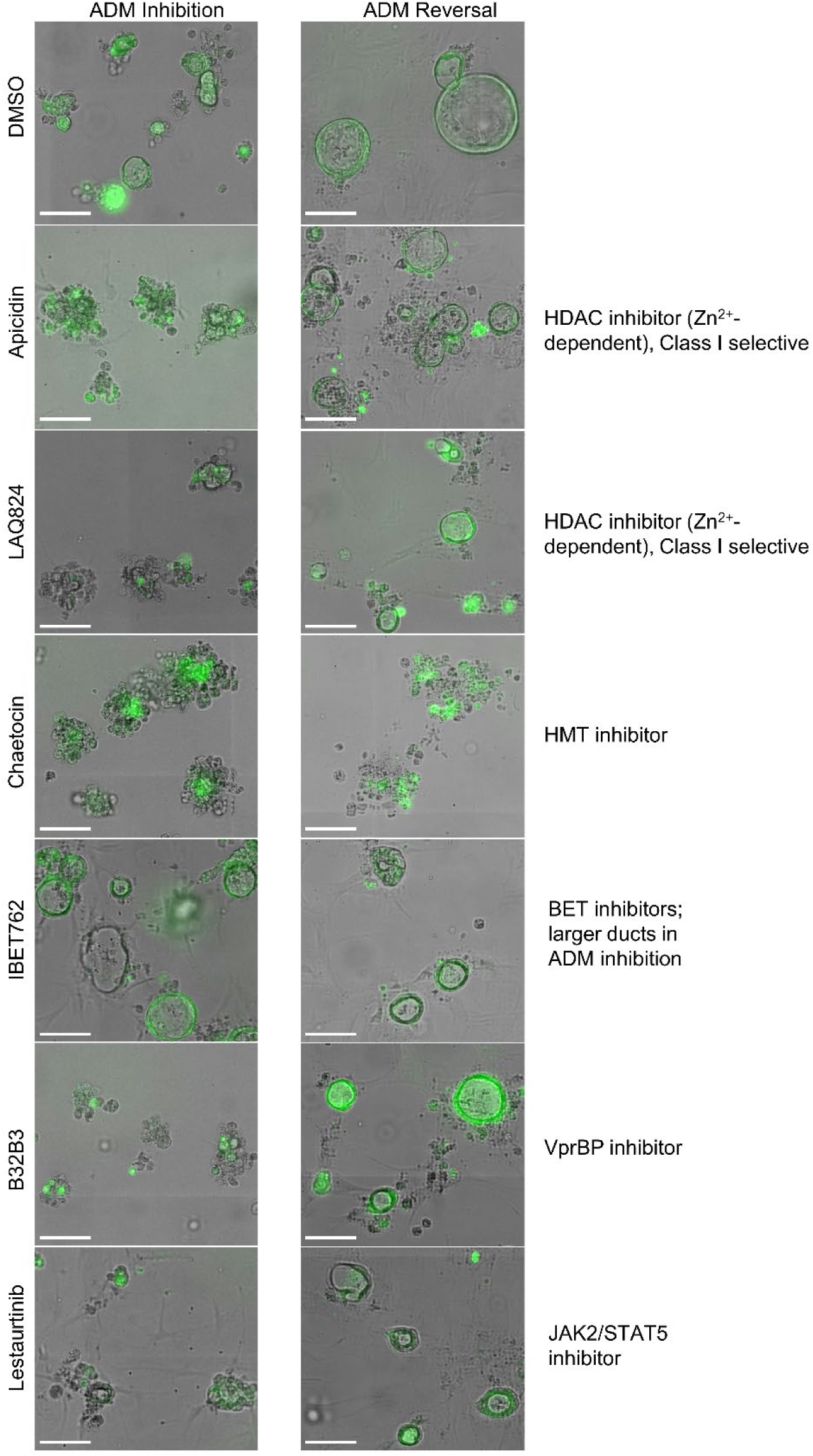
Cellular morphology of organoids following epigenetic small-molecule modulator library treatment in ADM inhibition (wildtype mice) or reversal (Cre mice) assays. Representative images of observed range of organoid morphology changes after treatment with epigenetic small-molecule modulator library compounds at 1 µM for 72 h in ADM inhibition (left) or ADM reversal (right). Size markers represent 100 µm.

### 3.4 Concentration-effect analysis for selected compounds

Based on our observations from the initial screens, a small number of compounds were selected for further validation (summarized in Table 1). The four bromodomain inhibitors IBET151, IBET762, (+)-JQ1 and PFI-1, that showed pronounced enlargement of formed ducts compared to those observed in the vehicle control were selected for further verification by concentration-effect analysis (Figure 8). The compounds LAQ824 (dacinostat, class I/II HDAC inhibitor) and lestaurtinib (tyrosine kinase inhibitor), showed only pronounced reduction in duct size (Figure 8), but not reversal to acinar morphology, and was therefore selected for follow up to see if reversal would be achieved at slightly higher concentrations. Also selected for concentration-effect analysis were apicidin and chaetocin as they were the prioritized compounds in both the inhibition and reversal screens.

The concentration-effect analysis was performed in wildtype (inhibition) and Cre (reversal) mouse organoid using concentrations ranging from 32 nM to 10 µM. Since apicidin and the positive control (trichostatin A) are both Zn^2+^-dependent HDAC inhibitors with differential selectivity profile (apicidin: class I; TSA: class I/II) and to possibly elucidate mode of action, we included in the follow up studies two class I-selective HDAC inhibitors: FK228, an FDA-approved anticancer drug derived from terrestrial bacteria, and largazole, a preclinical stage marine natural product [42] (Figure 9). Both of these compounds are class I HDAC inhibitors predominantly targeting HDACs 1, 2, and 3 and were not part of the ESL [42]. In terms of ADM reversal, FK228 was the most efficacious with low nM IC_50_, followed by apicidin, chaetocin and LAQ824; B32B3 was ineffective at reversing ADM (Figure 9). Of the six compounds selected for ADM inhibition, FK228 was the most potent with an IC_50_ of 16 nM (Figure 9). This was followed by apicidin, chaetocin, LAQ824 and B32B3 which were equipotent at IC_50_ ranging from ∼ 0.5 to 1 µM (Figure 9).

**Figure 9.**
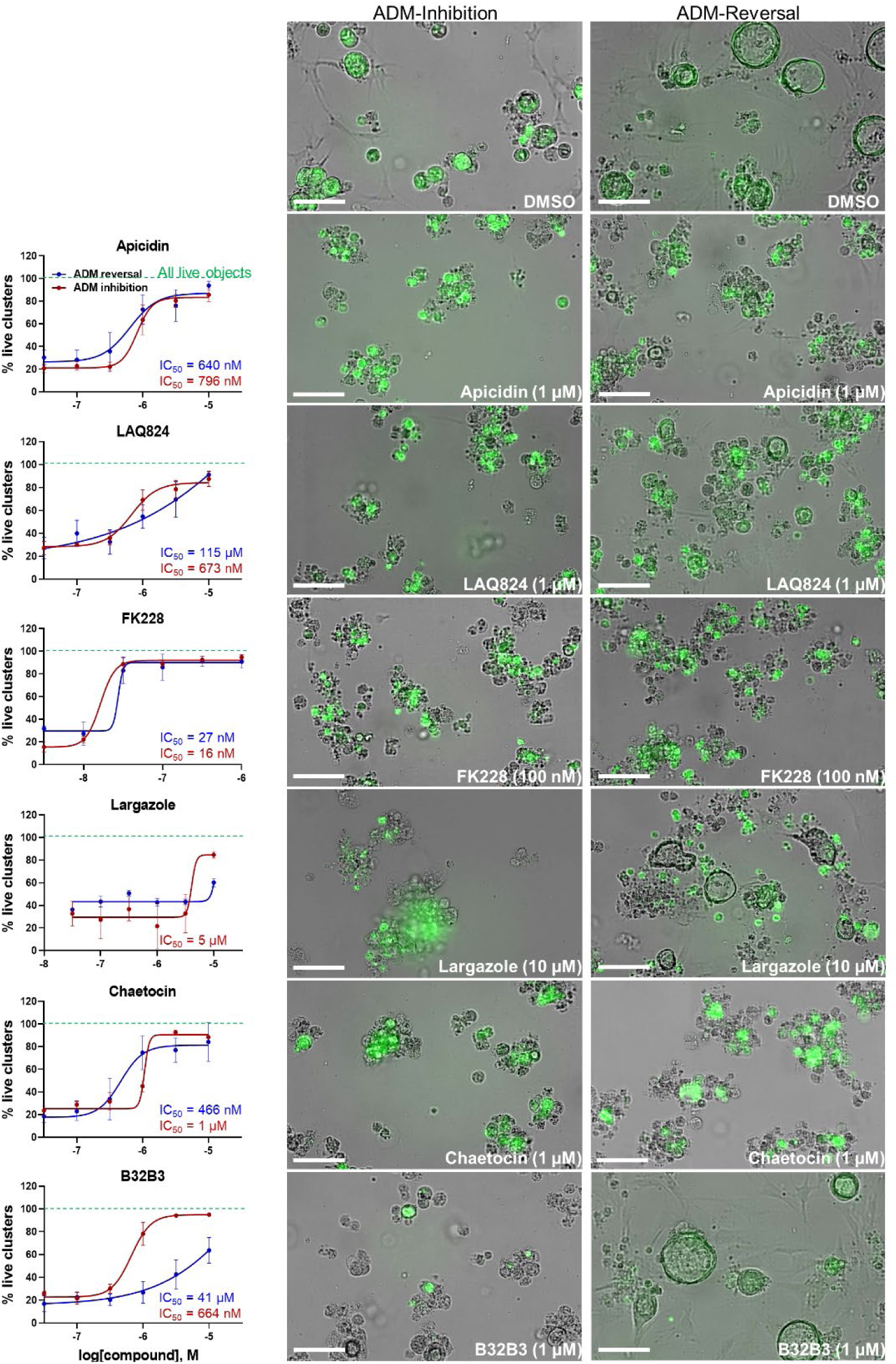
Validation of selected hits in dose response at 72 h post treatment. (Left) Percent live clusters from all live objects in organoids treated with dose response of select hits in ADM inhibition (wildtype mice) and ADM reversal (Cre mice) assay modes. IC_50_ values were calculated using GraphPad Prism software. (Right) Representative images of morphological changes observed with hit compounds at the lowest dose which caused morphologically distinguishable effects in ADM inhibition and ADM reversal assay modes. Size markers represent 100 µm.

Chaetocin was the only HMTase inhibitor compound from the library that showed similar effects in both the ADM inhibition (IC_50_ ∼ 1 µM) and the ADM reversal (IC_50_ ∼ 0.5 µM) assays. Compounds that did not validate with the initially observed results in either or both assay modes are shown in Figure S5. From the BET inhibitors selected for validation due to the unique effects, IBET151 (at 10 μM - 100 nM), IBET762 (at 10 μM – 32 nM) and (+) JQ1 (at 10 μM – 1 μM) showed enlargement of ducts in the ADM-inhibition assay, but no clear reversal in the ADM-reversal assay, and therefore we did not continue further analysis with these compounds. PFI-1 (the fourth BET inhibitor) did not show any effects in either mode during validation.

Although largazole has been previously shown to have similar potency to FK228 in enzyme and cell based (proliferation) assays in colorectal cancer HCT116 cells [42], it was less efficacious at both inhibiting and reversing ADM compared to FK228 (Figure 9). This may be due to its lower stability in the extracellular matrix where it can be hydrolyzed before entry into the cells and then would not reach the target [42; 43]. Largazole is a thioester prodrug that undergoes protein-assisted hydrolysis to liberate the active species, largazole thiol [42], while the disulfide FK228 is reductively activated in the cell, mediated by glutathione [44]. We have previously shown that the timing of prodrug activation can be modulated using disulfide homodimer and heterodimers [43]. Therefore, we performed an additional test in the ADM reversal mode assay with a largazole homodimer that was designed to have improved stability while also liberating two equivalents of active species, largazole thiol [43] (Figure 10). The homodimer showed similar efficacy during ADM reversal compared to apicidin, with an IC_50_ near 1 µM, but still 10-fold less activity compared to FK228 (IC_50_ near 0.1 µM) (Figures 9 and 10).

**Figure 10.**
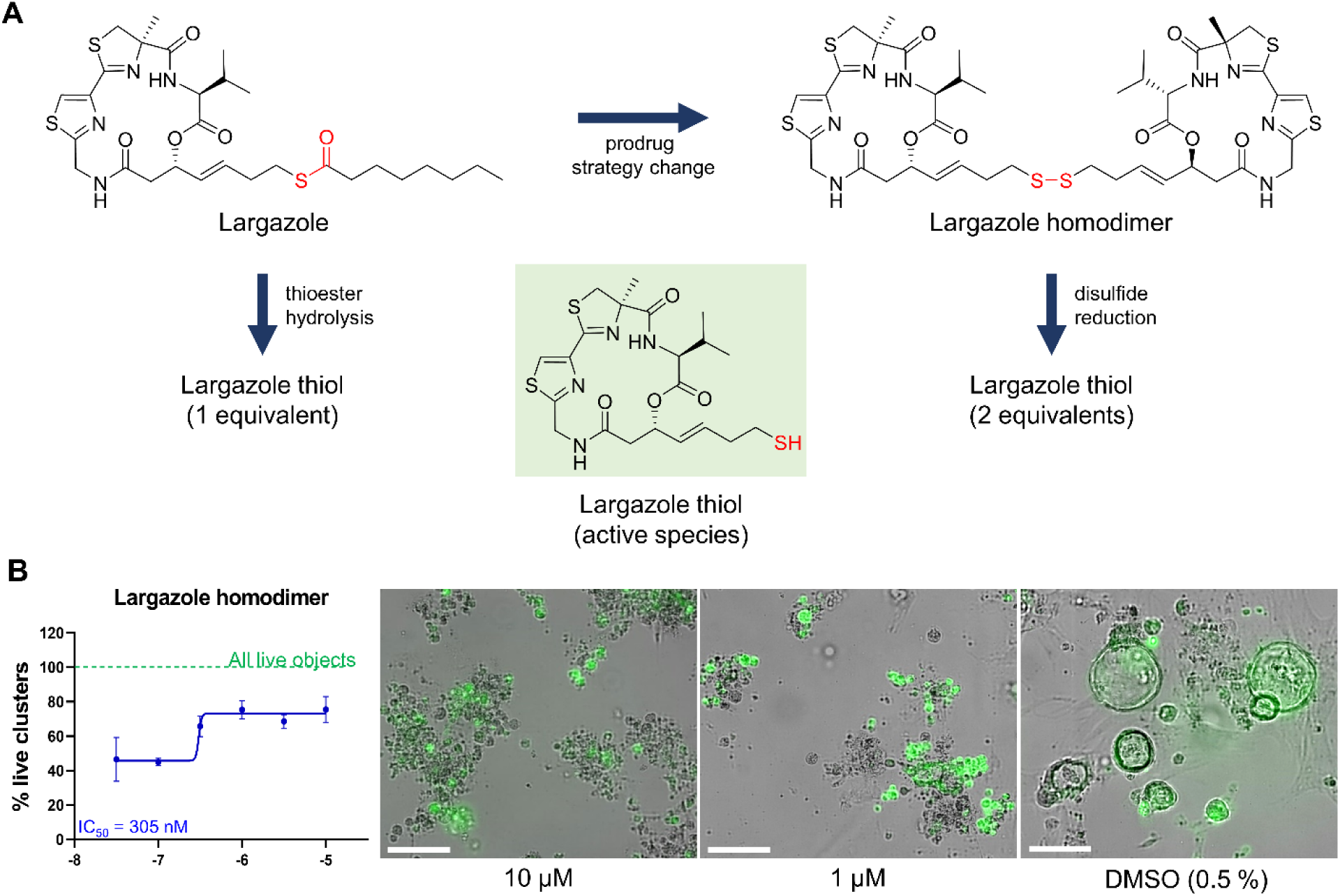
Testing of potency of largazole homodimer in ADM reversal mode (Cre mice organoids) at 72 h post treatment. (A) Structures of Largazole homodimer and justification for drug selection. Prodrug strategy was applied, which increases stability of the compound and also generates two equivalents of the active species, leading to increased potency compared to the parent compound. (B) Dose response study of largazole homodimer in ADM reversal assay mode. (Left) Percent live clusters from all live objects. IC_50_ value was calculated using GraphPad Prism 9 software based on quadruplicate experiments. (Right) Representative images of morphological changes observed with the homodimer at select doses in ADM reversal assay mode compared to vehicle control (0.5% DMSO). Scale bars represent 100 µm.

### 3.5 ADM reversal in KC mouse organoids

The validated compounds from both assays at 1 µM or lower were also tested for ADM reversal in the more clinically relevant *Kras*^G12D^-mutant mouse model (KC mice). These mice carry the *Kras^G12D^* gene mutation that is present in PDAC patients. Cultured KC organoids develop ADM faster than wild-type or Cre mouse organoids [45; 46]. In the KC mice, chaetocin concentrations of 1 and 3.2 µM were the most effective at reversing acinar morphology (Figure 11). In the vehicle control-treated organoids, the ducts progressed into obstructed ducts (cyst-like) that lacked a visible lumen [47]. Higher concentrations of the compounds apicidin, LAQ824, largazole homodimer and FK228 prevented the formation of these cyst-like structures compared to the DMSO control (Figure 11).

**Figure 11.**
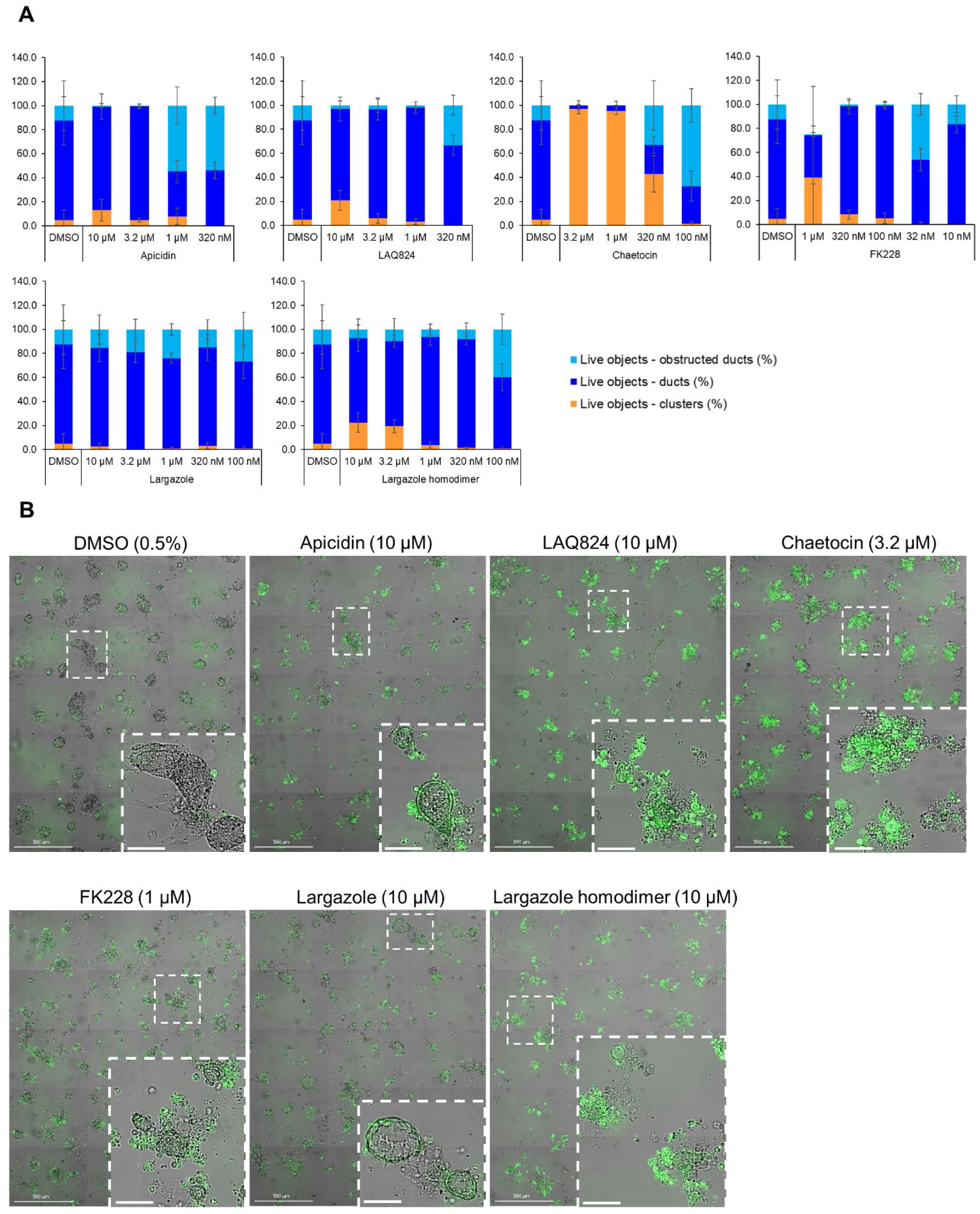
Effects of select validated compounds in KC mouse acini, ADM reversal assay. (A) Percent live clusters from all live objects in organoids treated with a small dose range of the compounds. (B) Representative images and enlarged area of the observed morphological effects. Size markers for full well images represent 500 µm and enlarged areas 100 µm.

qRT-PCR was used to validate that the morphological changes that occurred during ADM reversal of KC mouse organoids correlated to changes in acinar and ductal gene expression. Compared to DMSO control, all compounds tested produced predominate increased expression of the acinar genes *Amy2a*, *Cela1* and *Cpa2* with most occurring in a concentration-dependent fashion (Figure 12). Expression of the ductal genes were reduced by most of the treatments with largazole homodimer, FK228 and chaetocin demonstrating concentration-dependent reduction in *Krt19*, *Krt7* and *SOX9* expression (Figure 12). The ADM reversal index (ADMRI) was used as a measure of drug effect with the greater magnitude of ADMRI indicating more ADM reversal. The data show that for the 1 µM treatment FK228 was the most potent compound at reversing ADM while LAQ824 was most active at the 10 µM treatment (Table 2).

**Figure 12.**
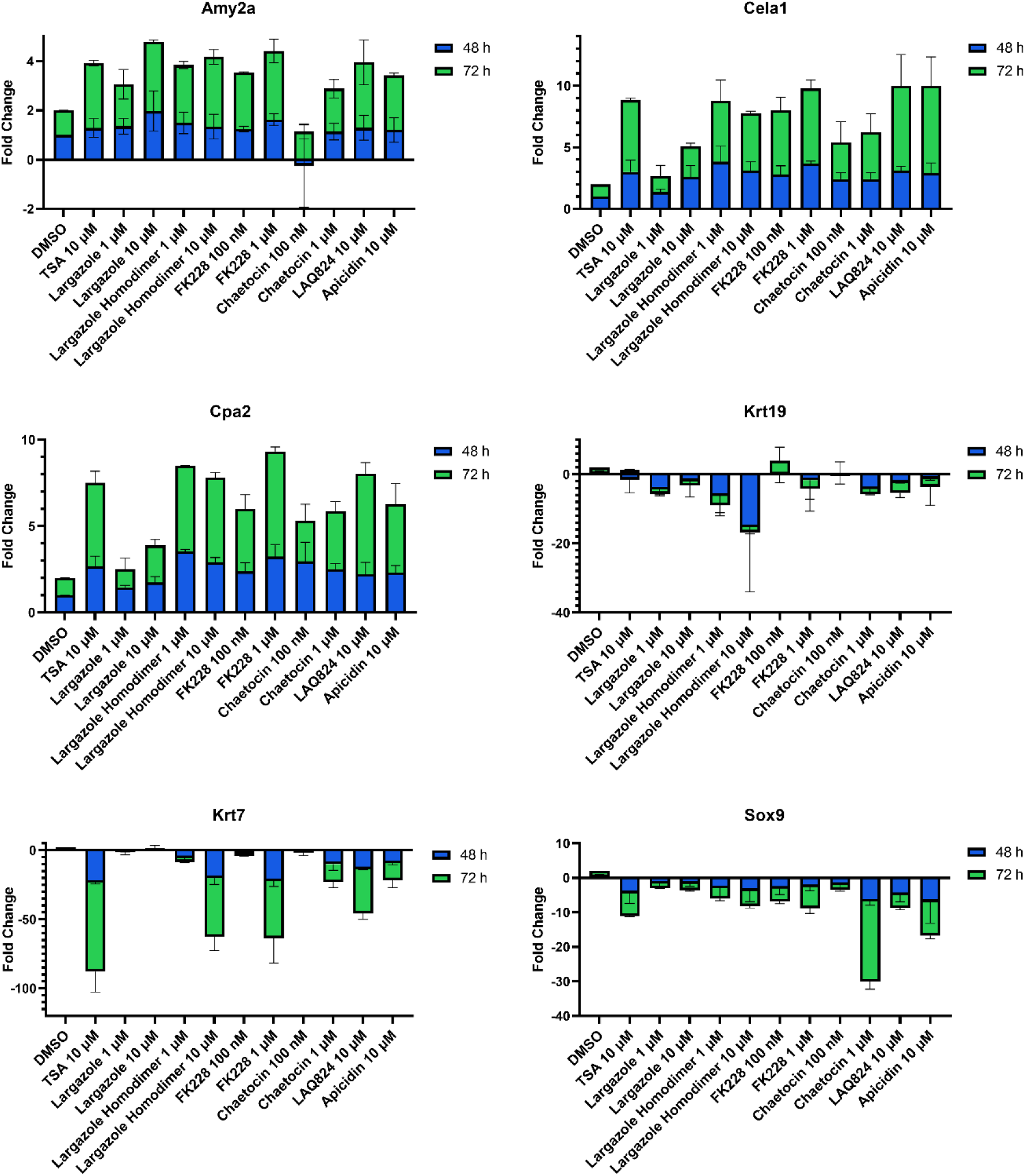
Validation of acinar and ductal gene expression following ADM reversal in KC organoids. KC mouse ADM organoids were treated with TSA, largazole, largazole homodimer, FK228, or chaetocin at varying concentrations for 2 or 3 days immediately following 2 days of ADM transdifferentiation. Cultures were collected following the 48 or 72 h treatments and gene expression was analyzed by qRT-PCR. Data are normalized to the 48 h DMSO control and are presented relative to 18S rRNA. Mean ± SD from two independent experiments.

RNA obtained from the ADM reversal experiments in KC mice for the most potent compounds at an IC_90_ dose were submitted for RNA sequencing to further validate if differences in gene expression accompanied the morphological changes and to see if pathway analysis support ADM reversal. The volcano plots of the drug-induced changes in gene expression show excellent correlations between acinar genes being upregulated and downregulation of ductal/PDAC genes upon reversal for largazole homodimer, FK228 and chaetocin (Figure S6). Pathway analysis showed that the top inhibited upstream regulator for all three compounds was angiotensinogen (AGT). Other PDAC-related pathways that were inhibited for all three compounds during ADM reversal include TGFB1 and TNF. The pathways consistently activated for all three compounds during ADM reversal include α-catenin and PPARGC1A (Figure 13).

**Figure 13.**
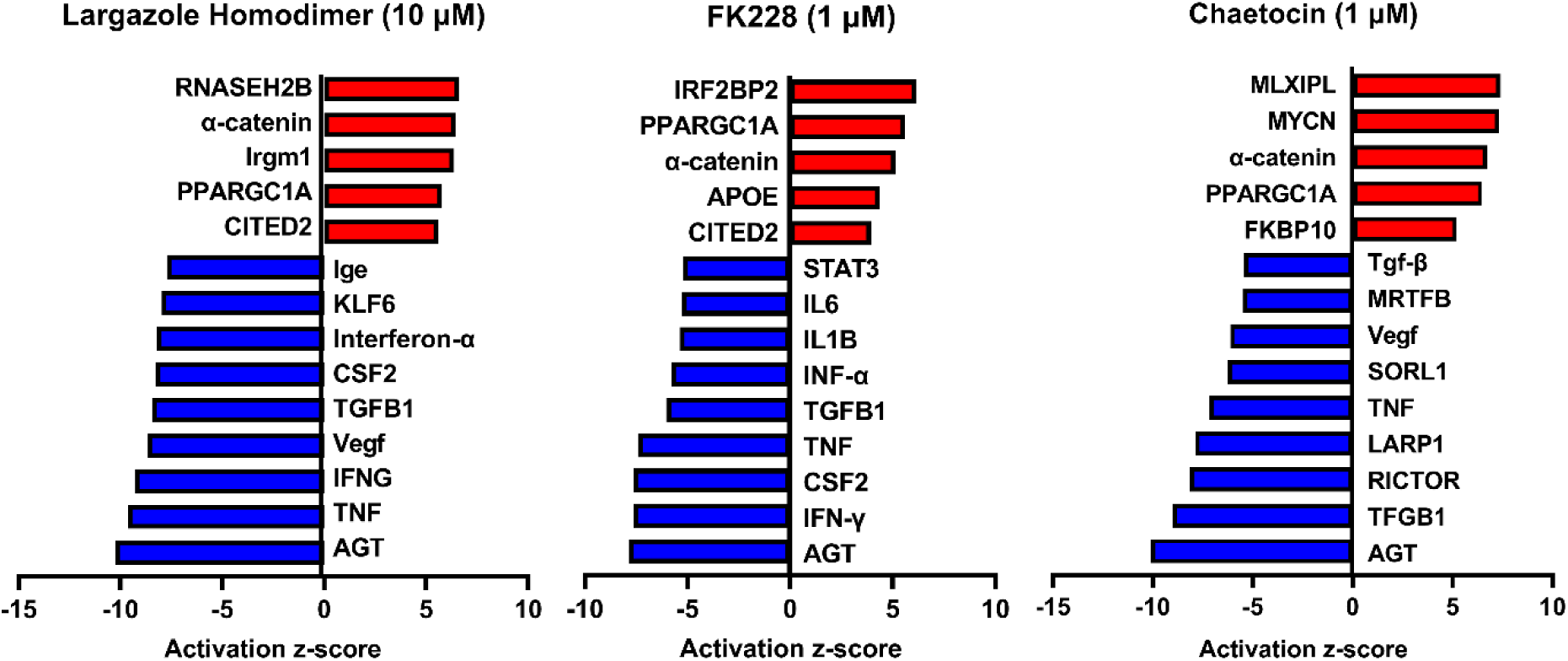
Ingenuity Pathway Analysis activation z-scores of regulated pathways from KC ADM reversal organoids. The RNA sequencing data from KC mice acinar organoids treated for ADM reversal with the compounds indicated were analyzed by Ingenuity Pathway Analysis (IPA) for the class of upstream regulators. The top upstream regulators are ranked by z-score with those regulators that are activated in red and those inhibited in blue.

## 4. Discussion

A novel phenotypic small-molecule screen based on organoid morphology was developed to discover epigenetic compounds that inhibit or reverse pancreatic ADM. Unlike anticancer agent screens that use cytotoxicity as an endpoint, quantification of acinar and ductal morphology using high-content imaging successfully identified compounds that inhibit or reverse ADM without inducing excessive cytotoxicity. For proof-of-concept, we utilized a focused library containing 144 epigenetic modulators with the goal of discovering effective compounds that act at the chromatin level. Using the HDAC inhibitor TSA as a positive control, reported in our prior work to inhibit and reverse ADM [24], our robust screen produced acceptable Z’ scores ranging from 0.56 to 0.61.

Our strategy was to first screen the libraries on wildtype (inhibition) and Cre (reversal) mouse organoids then evaluate the top candidates for ADM reversal using the KC mouse cultures. The inhibition screens in wildtype mouse organoids produced two compounds that passed our rigorous selection criteria. These compounds include the HMT inhibitor chaetocin and the Zn^2+^-dependent, class I HDAC inhibitor apicidin (Figures 4, 5). Likewise, apicidin and chaetocin were the only two compounds selected to reverse ADM in the Cre mouse organoid screen (Figures 6, 7). To probe the involvement of class I HDAC isoforms, two additional class I HDAC inhibitors that were not part of the ESL (FK228 and largazole), were also evaluated for ADM reversal and inhibition in organoids that contain wildtype *Kras*, thereby linking the activity to HDACs 1-3. To summarize our findings in organoids with wildtype *Kras*, FK228 was the most effective compound at inhibiting and reversing ADM (low nM IC_50_) whereas apicidin, LAQ824 and chaetocin required somewhat higher concentrations to exert a similar effect (Figures 9,10). We show that the stable homodimer of the natural product largazole [43; 48] has ADM reversal effects and that alteration of the prodrug type can be used to modulate the activity profile (Figure 10).

In the ADM inhibition and reversal screens in organoids containing wildtype *Kras*, certain compounds induced unique morphological changes that were identified following ADM inhibition or reversal. Of note, four different BET inhibitors (IBET762, IBET151, (+) JQ1 and PFI-1) formed enlarged ducts during ADM inhibition. (Figure 8 and Table 1). Bromodomain and extra-terminal (BET) proteins form complexes by binding to epigenetic marks such as acetylated lysine residues, reducing interaction between histones and DNA, thus increasing transcription. Nuclear expression of bromodomain containing 4 (BRD4), a BET protein important for enhancer-mediated transcription of cell-identity genes, was detected in normal acinar and duct cells and in the nucleus of acinar cells during ADM [49]. Combining tissue injury with shRNA knockdown of BRD4 in wildtype mice or mice harboring a *Kras* mutation showed that BRD4-suppressed cells effectively lost their acinar morphology and acquired ductal markers; the authors concluded that BRD4 impairs ADM during normal regeneration [50]. Thus, it is reasonable to presume that the enlarged ducts formed by BRD4 inhibition of ADM in our experiments resulted from their inability to dedifferentiate.

A total of 7 compounds (Table 2) were selected for their ability to reverse ADM in the clinically relevant KC mouse organoid model. While ADM in the context of mutant *Kras* is believed to be irreversible, we previously showed using both morphology and gene expression analysis that TSA reverted KC mouse pancreatic ducts to an acinar state [24]. We used the same endpoints here to establish ADM reversibility in KC mouse organoids by epigenetic modulating compounds. The morphology data presented in Figure 11 showed somewhat mixed results. While higher concentrations of chaetocin reversed nearly all of the ducts to acinar clusters, the other compounds (apicidin, LAQ824, FK228 and largazole homodimer) induced modest levels of ADM reversal by morphology even at concentrations as high as 10 uM (Figure 11). Examination of the gene expression data revealed that nearly all of the compounds increased acinar and reduced ductal gene expression to some extent (Figure 12). These findings were confirmed by presentation of the gene expression changes using the ADMRI which allowed us to rank the compound’s effectiveness (Table 2). These data confirm that compounds selected for their ability to reverse ADM in wildtype mouse organoids reverse ADM in organoids containing mutant *Kras*. Moreover, our screening approach successfully identified compounds that are more active at reversing ADM compared to the TSA positive control (Table 2, 10 µM treatments) and invoked the class I HDACs as major functionally relevant targets. Finally, our findings suggest that pharmacological re-expression of acinar genes and reduction in ductal genes may occur in cells undergoing ADM reversal that still morphologically resemble ducts.

It is noteworthy that in the KC mouse model, the top inhibited upstream regulator detected during ADM reversal (angiotensinogen, AGT) is the main precursor of angiotensin and part of the renin-angiotensin-aldosterone system (Figure 13). These data corroborate our findings of angiotensinogen as the most upregulated pathway during both human and mouse ADM [41]. Major components of the renin-angiotensin-aldosterone system (angiotensinogen and angiotensin receptors 1 and 2) were upregulated in rat pancreatic acinar cells during pancreatitis and treatment with the angiotensin receptor antagonist losartan inhibited the acinar digestion enzyme secretion [49]. Moreover, PDAC patients who were prescribed angiotensin converting enzyme inhibitors or angiotensin receptor blockers to treat their hypertension were associated with better clinical outcomes from gemcitabine monotherapy [51].

For organoids treated with the library at a final concentration of 1 μM, we identified and validated five compounds (apicidin, LAQ824, FK228, largazole and chaetocin) that induced complete or partial inhibition or reversal of ADM. Notably, of the five identified compounds, four (apicidin, LAQ824, FK228 and largazole) are Zn^2+^-dependent HDAC inhibitors, confirming the previously suggested importance of HDAC isoforms as possible targets for ADM modulation [24; 43; 48]. Our cumulative data indicate the importance of class I HDACs. FK228 and largazole specifically predominantly target HDACs 1, 2 and 3 and to a lesser extent HDAC8 [42]. The tested library contained a total of 34 Zn^2+^-dependent HDAC (class I/II/IV) inhibitors with broad or class-specific selectivity and 9 NAD^+^-dependent (class III) HDAC inhibitors, raising questions as to why the other class I HDAC inhibitors did not show similar effects as apicidin, LAQ824, FK228 and largazole. This could be due to the limitations of the current screen which used 1 μM as the primary screening concentration, selected based on a cytotoxicity screen of the ESL using cancer cell lines (data not shown). The 1 μM screening concentration ensured that over 80% of the tested compounds displayed less than 50% cytotoxicity, but it may fall below the IC_50_ of some of the compounds. Other possible reasons for the small amount of hits detected in this focused library screen could be a narrow target specificity, structure-specific physicochemical issues, or other differential downstream mechanisms involved that should be further explored. We have previously observed compound-specific effects of HDAC inhibitors in another small-molecule screen for epigenetic modulators of nuclear morphology [52].

## 5. Conclusion

In conclusion, we developed a novel phenotypic drug screen using organoid morphology as a readout to discover epigenetic regulator compounds with unique mechanism of action (i.e., ADM inhibition and reversal). Validation of the top hits (FK228, chaetocin, LAQ824 and largazole homodimer) in organoids derived from a clinically relevant KC mouse model confirmed that ADM can be reversed without inducing significant cytotoxicity even in the presence of mutant *Kras*. Our findings demonstrate a unique mechanism of action for epigenetic compounds and suggest that the phenotypic screen developed here may be applied to discover potential new treatments for PDAC.

### Data Availability Statement

All curated RNA-sequencing data sets were posted to the Gene Expression Omnibus (GEO) repository under accession number GES236292. Other data that support the findings of this study are available from the corresponding authors upon reasonable request.

### Ethics Statement

Transgenic animals were bred and studies performed at the University of Florida according to an approved IACUC protocol 202109058.

### Author Contributions

Kalina R. Atanasova carried all screening experiments and analysis, curated data and drafted manuscript. Corey M. Perkins plated organoids for all experiments, performed RNA extractions and RNA related data analysis, and manuscript editing. Ranjala Ratnayake designed and optimized screening, supervised all experiments, carried out manuscript writing and editing. Qi-Yin Chen performed chemical synthesis of largazoles. Jinmai Jiang performed RNA experiments and data analysis. Thomas D. Schmittgen provided project supervision, experimental design and manuscript writing. Hendrik Luesch provided project supervision, experimental design and manuscript editing.

### Funding

This research was in part supported by the NIH, National Cancer Institute Grants R01CA172310 (HL), Diversity Supplement R01CA172310-08S1 (CP), and R50CA211487 (RR), the Debbie and Sylvia DeSantis Chair Professorship (HL), College of Pharmacy PROSPER grant, and UF Health Cancer Center for screening support.

### Publisher’s Note

All claims expressed in this article are solely those of the authors and do not necessarily represent those of their affiliated organizations, or those of the publisher, the editors and the reviewers. Any product that may be evaluated in this article, or claim that may be made by its manufacturer, is not guaranteed or endorsed by the publisher.

## Acknowledgments

We acknowledge Lais Da Silva for assistance with organoid culture and organoid dispensing and James Matthews for pipeline development in pilot studies. We acknowledge Taylor Corcoran for support in preparing compound plates for screening.

