## Supplemental material for "Epigenetic Small-Molecule Screen for Inhibition and Reversal of Acinar Ductal Metaplasia in Mouse Pancreatic Organoids"

### Contents:

#### List of Figures:

**Figure S1**: Percent viability of organoids treated with ESL library compounds at 1 µM evaluated as percent live organoids from all organoids at 72 h post treatment in the ADM inhibition assay.

**Figure S2**: Percent viability of organoids treated with ESL library compounds at 1 µM evaluated as percent live organoids from all organoids at 72 h post treatment in the ADM reversal assay.

**Figure S3**: ADM-inhibition assay percent duct/cluster distribution (± standard deviation) of all objects at 72 h post treatment with the Cayman ESL, organized by groups of inhibitors.

**Figure S4**: ADM-reversal assay percent duct/cluster distribution (± standard deviation) of all objects at 72 h post treatment with the Cayman ESL, organized by groups of inhibitors.

**Figure S5**: Dose response effects on percent live clusters from all live objects of screen-selected compounds that did not validate at the screen-tested dose of 1 µM, or showed opposite effects (e.g. duct size enlargement) in ADM inhibition and ADM reversal assay modes.

**Figure S6:** Volcano plots of genes previously identified as those associated with pancreatic acinar/ductal phenotype or genes associated with onset or progression of PDAC.

**Table S1.** Annotation of genes associated with the pancreatic acinar/ductal phenotype or with the onset or progression of PDAC

**Supplementary Materials and Methods:** RNA sequencing and pathway enrichment analysis

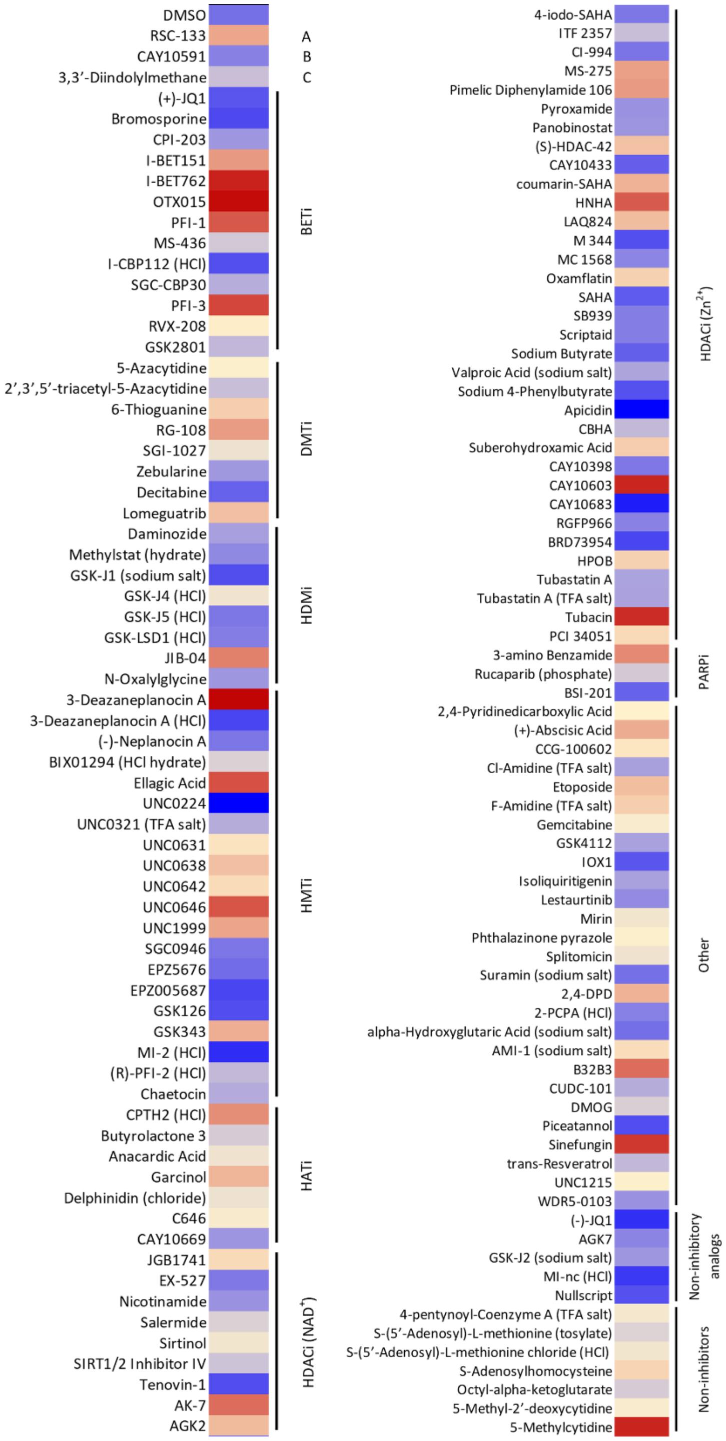


| Total ESL compounds | 144 |
|---|---|
| ≤50% viability at 1 µM | 47 |
| >50% viability at 1 µM | 97 |

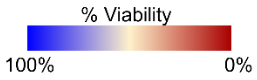


**Figure S1. Percent viability of organoids treated with ESL library compounds at 1 µM final concentration evaluated as percent live organoids from all organoids at 72 h post treatment in the ADM-inhibition assay.** Viability of organoids was evaluated using Calcein AM viability stain. Abbreviations: BETi = bromodomain and extra-terminal domain inhibitors; DMTi = DNA methyltransferase inhibitors; HDMi = histone demethylase inhibitors; HMTi = histone methyltransferase inhibitors; HATi = histone acetyltransferase inhibitors; HDACi = histone deacetylase inhibitors; PARPi = poly-ADP ribose polymerase inhibitors. A = DMT and HAT inhibitor; B = HDAC activator ($NAD^+$ dependent); C = HMT and DMT inhibitor. HCl = hydrochloride; TFA salt = trifluoroacetic acid salt.

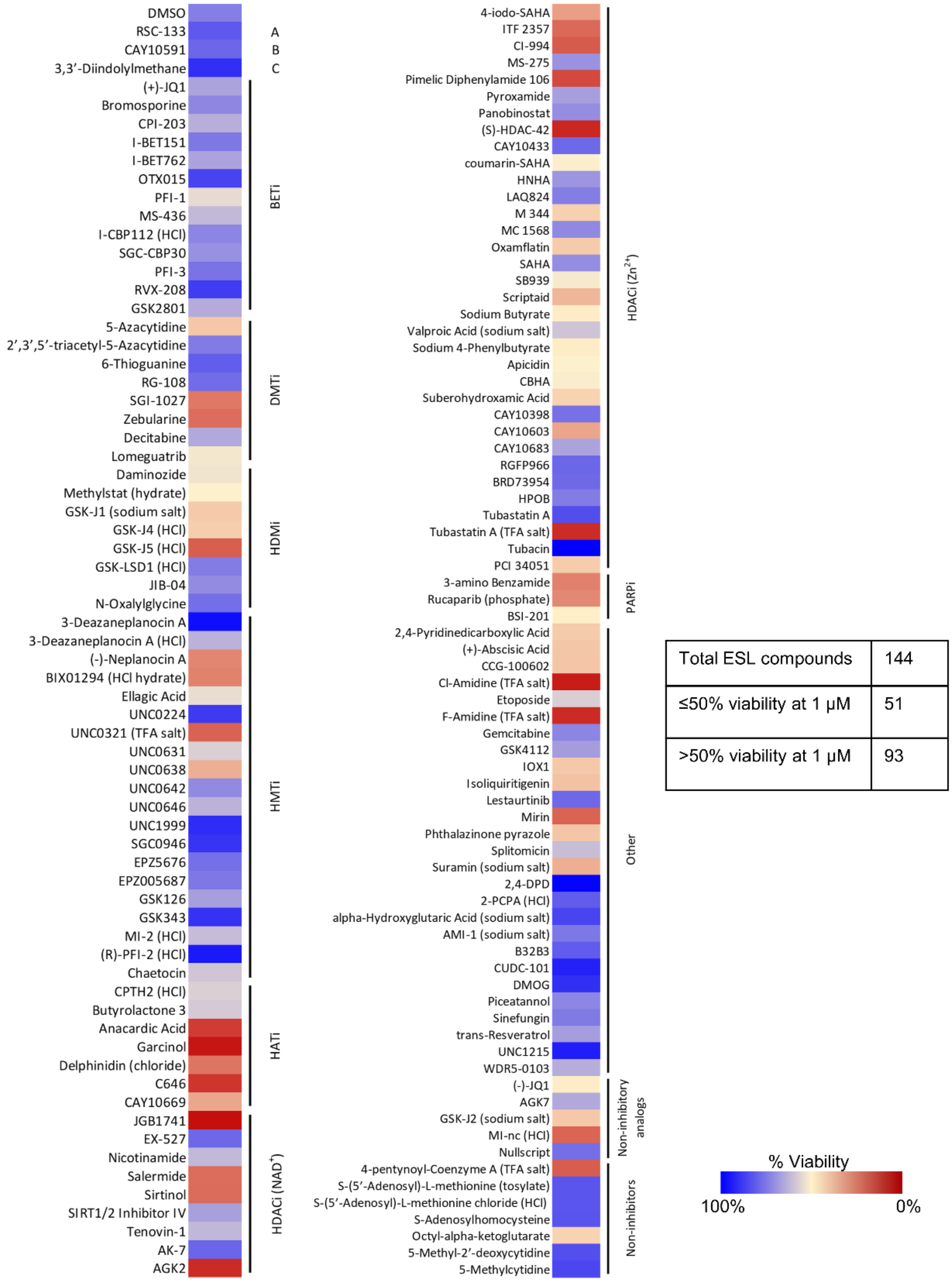


| Total ESL compounds | 144 |
|---|---|
| ≤50% viability at 1 μM | 51 |
| >50% viability at 1 μM | 93 |

**Figure S2**. **Percent viability of organoids treated with ESL library compounds at 1 μM final concentration evaluated as percent live organoids from all organoids at 72 h post treatment in the ADM-reversal assay.** Viability of organoids was evaluated using Calcein AM viability stain. Abbreviations: BETi = bromodomain and extra-terminal domain inhibitors; DMTi = DNA methyltransferase inhibitors; HDMi = histone demethylase inhibitors; HMTi = histone methyltransferase inhibitors; HATi = histone acetyltransferase inhibitors; HDACi = histone deacetylase inhibitors; PARPi = poly-ADP ribose polymerase inhibitors. A = DMT and HAT inhibitor; B = HDAC activator ($NAD^+$ dependent); C = HMT and DMT inhibitor. HCl = hydrochloride; TFA salt = trifluoroacetic acid salt.

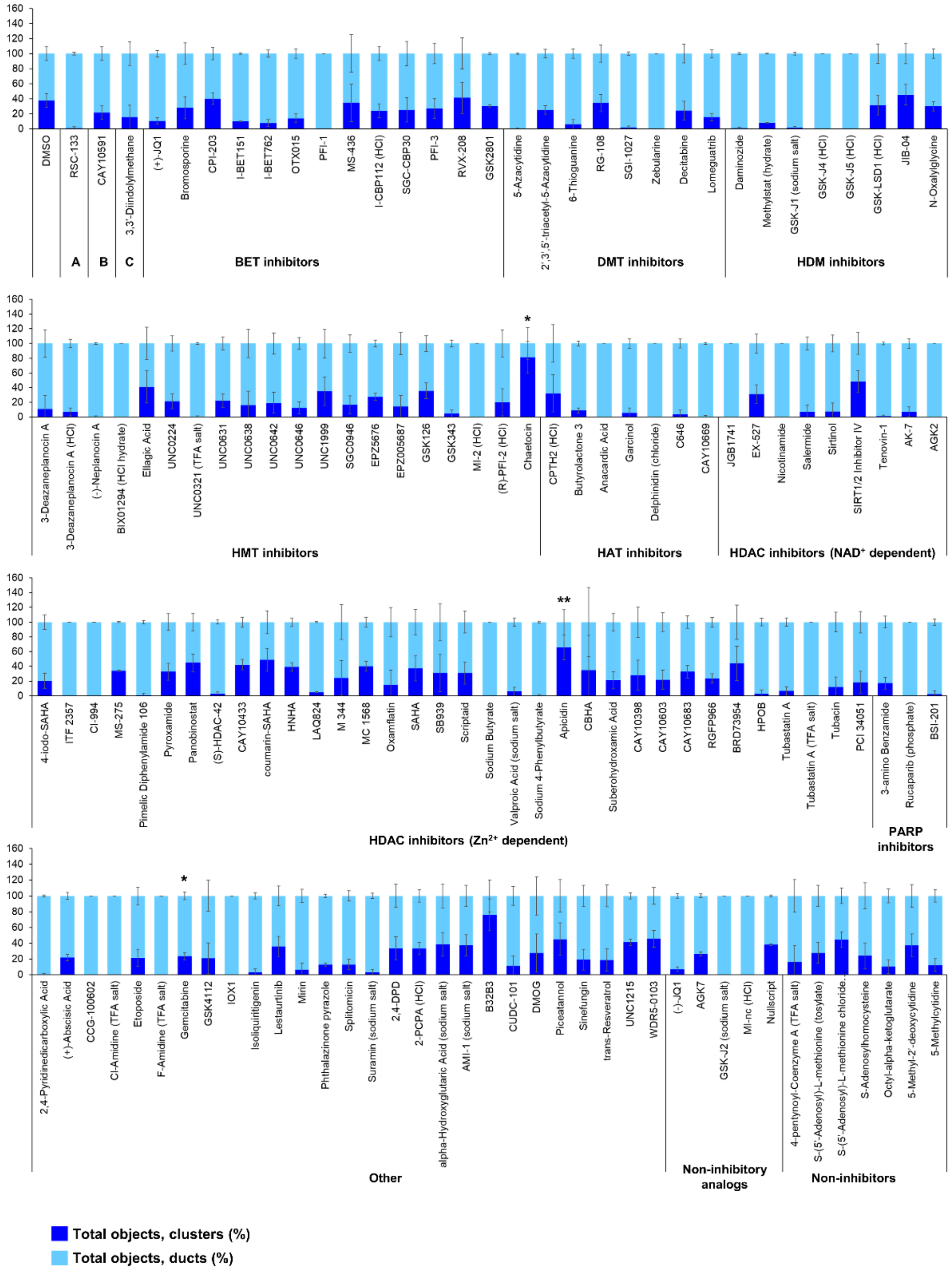


**Figure S3**. **ADM-inhibition assay percent duct/cluster distribution (± standard deviation) of all objects at 72 h post treatment with the Cayman ESL, organized by groups of inhibitors.** Abbreviations: BET = bromodomain and extra-terminal domain inhibitors; DMT = DNA methyltransferase inhibitors; HDM = histone demethylase inhibitors; HMT = histone methyltransferase inhibitors; HAT = histone acetyltransferase inhibitors; HDAC = histone deacetylase inhibitors; PARP = poly-ADP ribose polymerase inhibitors. A = DMT and HAT inhibitor; B = HDAC activator (NAD$^+$ dependent); C = HMT and DMT inhibitor; HCl = hydrochloride; TFA salt = trifluoroacetic acid salt. P-values were calculated using two-tailed Student’s t-test with unequal variances. Significance was accepted at P≤0.05 only when averages of clusters are higher than the respective vehicle control. * P ≤0.05 – 0.01; ** P <0.01 – 0.001; *** P <0.001.

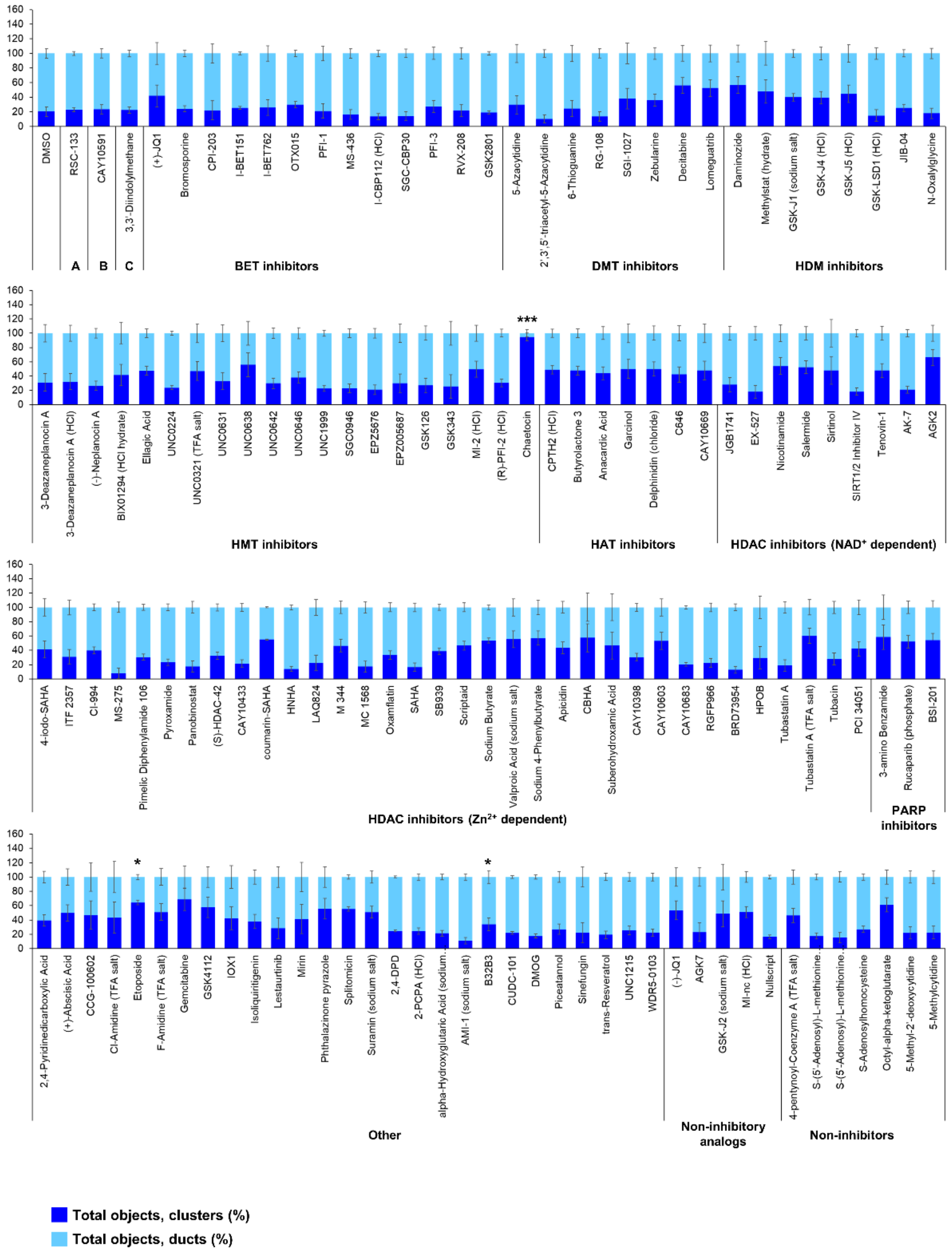


**Figure S4**. **ADM-reversal assay percent duct/cluster distribution (± standard deviation) of all objects at 72 h post treatment with the Cayman ESL, organized by groups of inhibitors.** Abbreviations: BET = bromodomain and extra-terminal domain inhibitors; DMT = DNA methyltransferase inhibitors; HDM = histone demethylase inhibitors; HMT = histone methyltransferase inhibitors; HAT = histone acetyltransferase inhibitors; HDAC = histone deacetylase inhibitors; PARP = poly-ADP ribose polymerase inhibitors. A = DMT and HAT inhibitor; B = HDAC activator ($NAD^+$ dependent); C = HMT and DMT inhibitor; HCl = hydrochloride; TFA salt = trifluoroacetic acid salt. P-values were calculated using two-tailed Student's t-test with unequal variances. Significance was accepted at $P \leq 0.05$ only when averages of clusters are higher than the respective vehicle control. * $P \leq 0.05 – 0.01$; ** $P < 0.01 – 0.001$; *** $P < 0.001$.

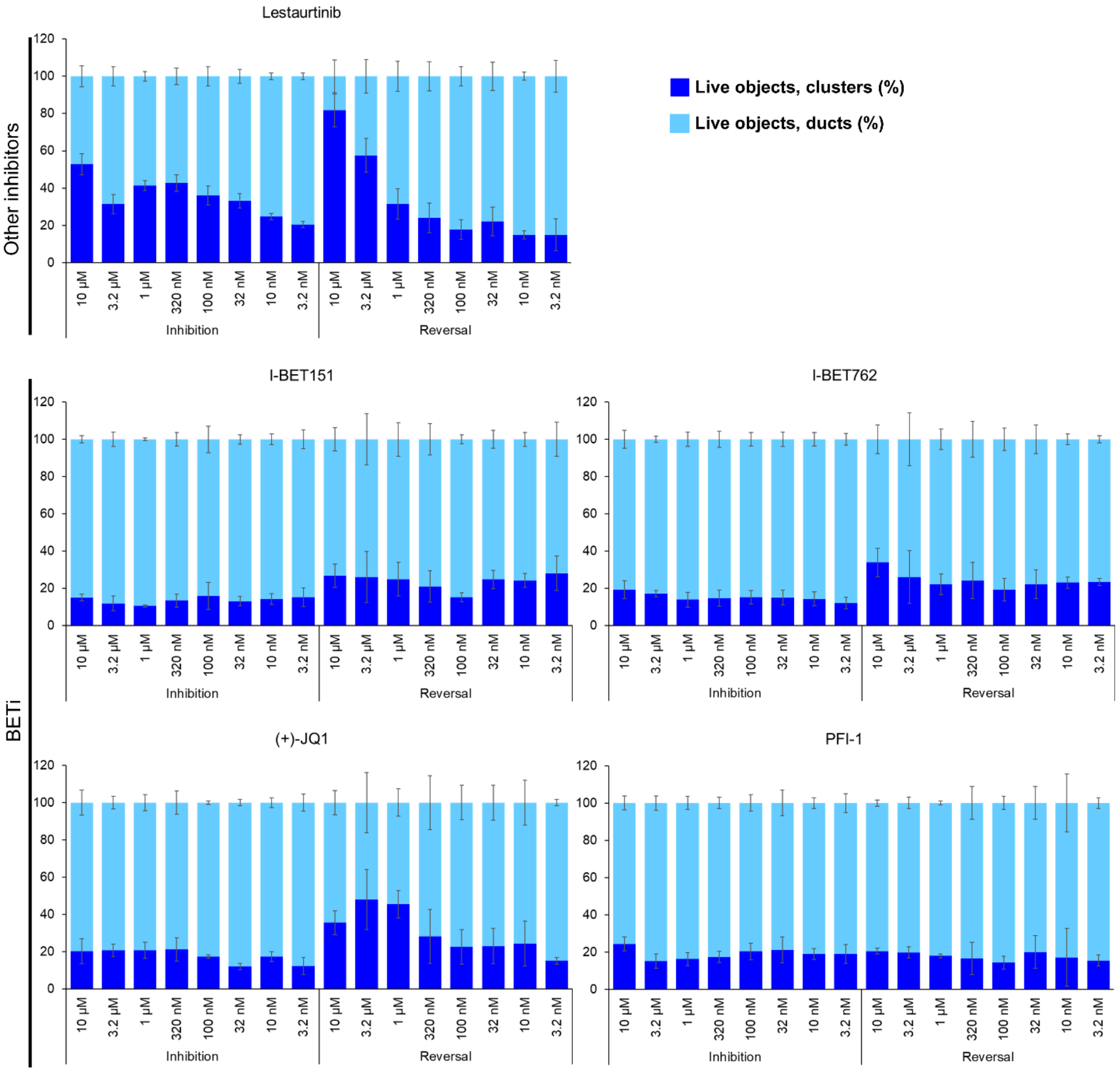


**Figure S5**. **Dose-response effects on percent live clusters from all live objects of screen-selected compounds that did not validate at the screen-tested dose of 1 µM, or showed opposite effects (e.g. duct size enlargement) in ADM-inhibition and ADM-reversal assay modes.**

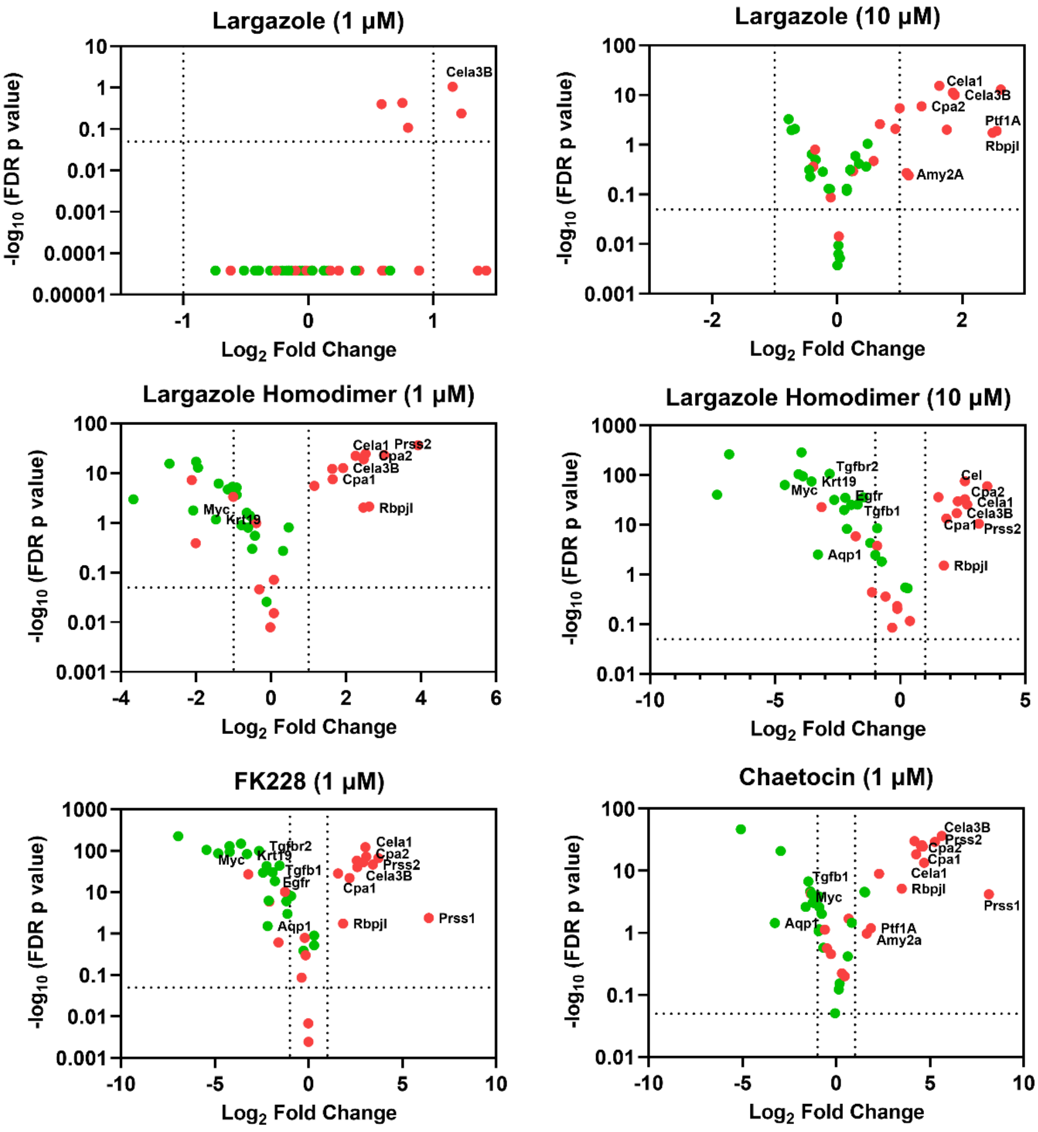


**Figure S6: Volcano plots of genes previously identified as those associated with pancreatic acinar/ductal phenotype or genes associated with onset or progression of PDAC.** RNA isolated from cultures of KC mouse acini that underwent ADM reversal by the compounds listed was sequenced. Fifty known acinar, ductal, and PDAC associated genes were selected and annotated based on the literature review (Supplemental Table 1). Volcano plots were constructed with the acinar (red) and ductal/PDAC (green) gene expression data indicated.

**Table S1.** Annotation of genes associated with the pancreatic acinar/ductal phenotype or with the onset or progression of PDAC

| Gene | tumor/acinar_genes | Gene | Ductal/acinar_sc |
|---|---|---|---|
| **EGFR** | tumor_genes | **ALB** | acinar_genes |
| **KRT19** | tumor_genes | **REG1B** | acinar_genes |
| **CD44** | tumor_genes | **PRSS1** | acinar_genes |
| **TGFB1** | tumor_genes | **CEL** | acinar_genes |
| **SMAD3** | tumor_genes | **PNLIP** | acinar_genes |
| **MYC** | tumor_genes | **CTRB2** | acinar_genes |
| **PPARG** | tumor_genes | **CPA2** | acinar_genes |
| **RUNX1** | tumor_genes | **ALDH1A3** | ductal_genes |
| **HES1** | tumor_genes | **CFTR** | ductal_genes |
| **MMP14** | tumor_genes | **AQP1** | ductal_genes |
| **ITGB1** | tumor_genes | **DEFB1** | ductal_genes |
| **IGF1R** | tumor_genes | **KRT19** | ductal_genes |
| **ERBB2** | tumor_genes | **SPP1** | ductal_genes |
| **AKT3** | tumor_genes | **TSPAN8** | ductal_genes |
| **TGFBR2** | tumor_genes | | |
| **NOTCH2** | tumor_genes | | |
| **AMY2A** | acinar_genes | | |
| **CPA1** | acinar_genes | | |
| **NR5A2** | acinar_genes | | |
| **EPCAM** | acinar_genes | | |
| **CEL** | acinar_genes | | |
| **PRSS2** | acinar_genes | | |
| **CELA3B** | acinar_genes | | |
| **PTF1A** | acinar_genes | | |
| **PNLIP** | acinar_genes | | |
| **PDE3B** | acinar_genes | | |
| **CELA1** | acinar_genes | | |
| **BHLHA15** | acinar_genes | | |
| **RBPJL** | acinar_genes | | |
| **IGF1** | acinar_genes | | |
| **BMP7** | acinar_genes | | |
| **MIST1** | acinar_genes | | |
| **E47** | acinar_genes | | |
| **TFE2** | acinar_genes | | |
| **DICER1** | acinar_genes | | |
| **PAF1** | acinar_genes | | |

**Supplementary materials and methods:**

RNA sequencing and pathway enrichment analysis: Illumina RNA library construction was performed at the Interdisciplinary Center for Biotechnology Research (ICBR) Gene Expression Core, University of Florida (UF). RNA quantitation was done on a NanoDrop Spectrophotometer (NanoDrop Technologies, Inc.), and sample quality was assessed using the Agilent 2100 Bioanalyzer (Agilent Technologies, Inc). SMART-Seq V4 ultra low input RNA kit will bewere used for RNAseq library construction according to the user manual. Briefly, 1st strand cDNA is primed by a 3' SMART-Seq CDS Primer II A, and SMART-Seq v4 Oligonucleotide was used for template switching at the 5' end of the transcript. Then cDNA was amplified by PCR Primer II A for 10 cycles. Finally, Illumina sequencing libraries were generated with 125 pg of cDNA using Illumina Nextera DNA Sample Preparation Kit (Cat#: FC-131-1024) according to manufacturer's instructions. Briefly, 125 pg of cDNA was tagmented and then adapter sequences added onto template cDNA by PCR amplification. Libraries were quantitated by Bioanalyzer and qPCR (Kapa Biosystems, catalog number: KK4824). Finally, the libraries were pooled equal molar concentration and sequenced by Illumina NovaSeq6000 Sequencing. Briefly, normalized libraries were submitted to the "Free Adapter Blocking Reagent" protocol (FAB, Cat# 20024145) in order to minimize the presence of adaptor-dimers and index hopping rates. The library pool was diluted to 0.8 nM and sequenced on one S4 flow cell lane (2x150 cycles) of the Illumina NovaSeq6000. The instrument's computer utilized the NovaSeq Control Software v1.6. Cluster and SBS consumables were v1.5. The final loading concentration of the library was 120 pM with 1% PhiX spike-in control. One lane generates 2.5-3 billion paired-end reads (~950Gb) with an average Q30%>= 92.5% and Cluster PF= 85.4%. FastQ files were generated using the BCL2fastQ function in the Illumina BaseSpace portal. Sequencing was performed at the ICBR NextGen Sequencing (https://biotech.ufl.edu/next-gen-dna/, RRID:SCR_019152).
